# Cross-species evidence for a developmental origin of adult hypersomnia with loss of synaptic adhesion molecules *beat-Ia/CADM2*

**DOI:** 10.1101/2024.09.25.615048

**Authors:** Kyla Mace, Amber Zimmerman, Alessandra Chesi, Fusun Doldur-Balli, Hayle Kim, Erika Almeraya Del Valle, Allan I. Pack, Struan F.A. Grant, Matthew S. Kayser

## Abstract

Idiopathic hypersomnia (IH) is a poorly-understood sleep disorder characterized by excessive daytime sleepiness despite normal nighttime sleep. Combining human genomics with behavioral and mechanistic studies in fish and flies, we uncover a role for *beat-Ia/CADM2*, synaptic adhesion molecules of the immunoglobulin superfamily, in excessive sleepiness. Neuronal knockdown of Drosophila *beat-Ia* results in sleepy flies and loss of the vertebrate ortholog of *beat-Ia*, *CADM2*, results in sleepy fish. We delineate a developmental function for *beat-Ia* in synaptic elaboration of neuropeptide F (NPF) neurites projecting to the suboesophageal zone (SEZ) of the fly brain. Brain connectome and experimental evidence demonstrate these NPF outputs synapse onto a subpopulation of SEZ GABAergic neurons to stabilize arousal. NPF is the Drosophila homolog of vertebrate neuropeptide Y (NPY), and an NPY receptor agonist restores sleep to normal levels in zebrafish lacking *CADM2*. These findings point towards NPY modulation as a treatment target for human hypersomnia.

## Introduction

Disrupted sleep is associated with diverse negative physiological consequences ^1–3^. Over the past few decades, work in animal models and humans has led to new insights into genetic regulation of sleep ^4–7^, and more recent efforts have leveraged human genome wide association studies (GWAS) to examine the genetics of numerous sleep disorders ^8–10^. However, in contrast to disorders like sleep apnea and insomnia, few studies have focused on disorders of unexplained excessive sleepiness. Idiopathic hypersomnia (IH) is characterized by excessive daytime sleepiness despite normal nighttime sleep, often accompanied by difficulty awakening (sleep inertia), unrefreshing sleep, and cognitive impairment (sleep drunkenness) ^11–13^. In contrast to type 1 narcolepsy, a hypersomnolence disease whose pathophysiology is well-understood, the etiology of IH is completely opaque. This lack of knowledge contributes to underdiagnosis of IH and poor management of its symptoms ^14^. Treatment strategies for this disorder have been hampered by a dearth of mechanistic insights. IH may be caused heterogenous etiologies, with hypothesized pathophysiologies relating to the immune system ^14^, circadian clock ^15,16^, and GABAergic dysfunction ^17^. Yet, mechanistic evaluation of these or other hypotheses remains limited.

Regardless of underlying cause, IH has a heritable component: twin studies show that daytime sleepiness is between 37-48% heritable and family history of excessive sleepiness is present in ∼1/3 of IH patients ^11,18–20^. Recent GWAS focused on sleep and circadian rhythms have provided new insights into sleep in health and disease ^3^. Discerning the biological impact of implicated loci remains a challenge, as little is known of their functional effects, due in part to the often-erroneous assumption that the nearest gene is the causal effector gene ^21^. Non-coding regions of the genome contain important regulatory elements and genetic variants that disrupt those elements can confer susceptibility to complex disease; such regulatory elements can be at substantial distance from the actual effector gene. “Variant-to-gene” mapping approaches have been applied to identify functional effects of variants emerging from insomnia GWAS, followed by validation in animal models ^8^. However, even these approaches can fail to fully account for potential genes impacted by a given variant. For example, a regulatory variant is more likely to affect expression in cis of a gene within its topologically associated domain (TAD) rather than elsewhere in the genome ^22^. A TAD is defined by Hi-C approaches as a self-interacting genomic region ^22^. As such, DNA regions within a TAD physically interact with each other more frequently than with regions outside the TAD ^23,24^. Prioritizing candidate genes within TADs has allowed for the identification of distal disease genes ^25^, but such a comprehensive approach has never been applied to sleep traits.

Here, combining human genomics with behavioral and mechanistic studies in fish and flies, we investigate molecular and genetic underpinnings of hypersomnia. Using a full TAD-wise analysis of genome wide significant loci associated with hypersomnia traits, we conducted a reverse-genetic screen in *Drosophila* to implicate candidate effector genes. This screen revealed a role for the synaptic adhesion molecule *beat-Ia* and its vertebrate ortholog, *CADM2*, in excessive sleepiness, coinciding with the napping and daytime sleepiness GWAS locus on human chromosome 3. Detailed analyses in flies support a developmental role for *beat-Ia* and its receptor, *side,* in coordinating synaptic elaboration of wake-promoting neuropeptide F (NPF) neurites in the suboesophageal zone (SEZ) of the fly brain. Loss of *beat-Ia* disrupts NPF synaptic input to GABAergic targets in the SEZ, resulting in impaired ability to maintain wakefulness. NPF is the *Drosophila* homolog of vertebrate neuropeptide Y (NPY) and we find that an NPY receptor agonist restores normal sleep/wake balance to zebrafish lacking *CADM2,* demonstrating how cross-species approaches may uncover novel treatment targets for human hypersomnia.

## Results

### Identification and RNAi screen of candidate hypersomnia-associated genes

To identify molecular and genetic factors contributing to excessive sleepiness, we conducted a reverse-genetic knockdown screen of hypersomnia-associated genes (**Fig. 1** for schematic). The most comprehensive GWAS for sleep reported to date are based on the UK Biobank (n=452,071) where 243 loci were identified ^26,26,27^. Using objective data about sleep in a subset of participants with 7-day accelerometry, 24 loci were initially associated with sleep propensity (excessive sleep duration, daytime napping, and daytime sleepiness), although 59 other loci were associated with sleep fragmentation, presumably secondarily driving daytime sleepiness and therefore less relevant to IH ^26–28^. Additionally, the first GWAS of IH patients (n=414) revealed 4 loci specifically associated with IH ^29^. With the goal of investigating the sleep propensity loci as exhaustively as possible, we identified all genes residing within the corresponding TADs harboring each GWAS signal from the 28 sleep propensity and hypersomnia loci identified. For this analysis, we used TAD coordinates from neural progenitor cells derived from the human H1 ESC line ^22^. This approach resulted in 26 TADs harboring 511 human genes, 274 of which are protein coding, coinciding with sleep propensity GWAS signals (**Fig. 1A, Supplemental Data 1**).

**Fig. 1.**
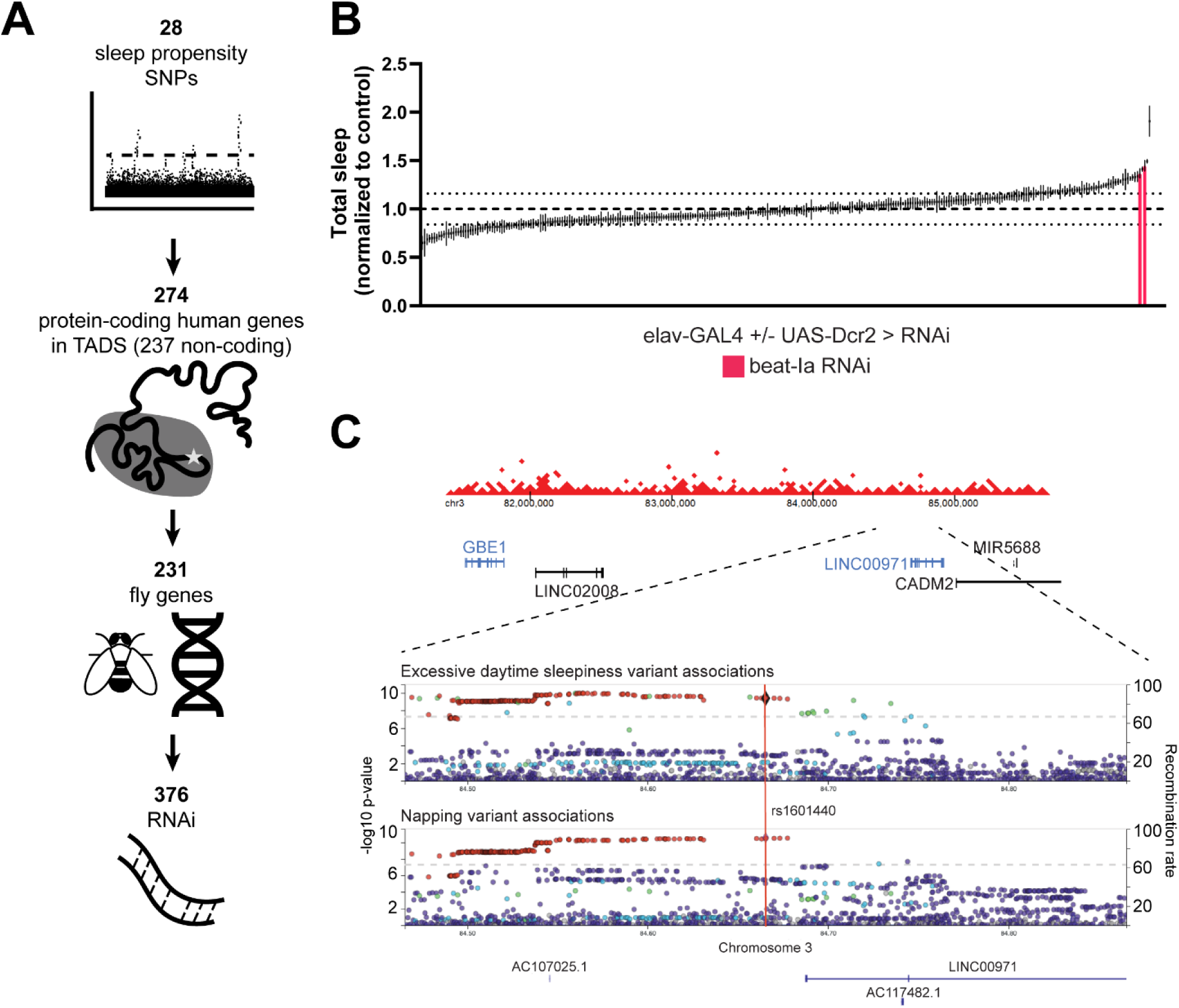
A reverse-genetic screen of hypersomnolence-associated variants identifies *beat-Ia* as a sleep regulatory molecule. (A) Schematic of gene candidate nomination and sleep screen in Drosophila. B) Total sleep across 24 hours of flies expressing RNAi driven by pan-neuronal elav-GAL4. UAS-Dcr2 was co-expressed for RNAi of the VALIUM1 or VALIUM10 vector to improve RNAi efficacy. Sleep duration for each experimental genotype was normalized to the control genotype of the experiment in which it was run. Lines indicate standard error of sleep measurements from 8-16 animals, except for 3 RNAi lines which have 3-7 animals each. Two RNAi against *beat-Ia* (magenta) increase total sleep duration beyond 2 standard deviations from the mean (indicated by dotted lines). (C) Top: Topologically associated domain containing the GWAS association signal for putative sleep propensity loci (in neural progenitor cells). TAD coordinates were retrieved from 3D Genome Browser (H1-NPCs) (http://3dgenome.fsm.northwestern.edu/index.html). Genes in black are on + strand, genes in blue are on – strand. Bottom: Locus zoom plots of the chromosome 3 region associated with napping and daytime sleepiness retrieved from the Sleep Disorder Knowledge Portal (https://sleep.hugeamp.org/). Lead SNP for napping rs1601440 is indicated by the orange line. Nearest gene is *LINC00971* (non-coding). Nearest protein coding gene is *CAMD2* (see TAD). LD (EUR) is indicated by color for sentinel SNP rs1601440.

Using DIOPT, an integrated ortholog prediction tool that combines numerous algorithms ^30^, we identified 231 fly orthologs for these human genes, and 376 publicly-available RNAi lines targeting the fly genes (**Fig. 1A**). We then screened sleep parameters in flies with expression of individual RNAi under control of the pan-neuronal driver, elav-GAL4 (**Fig. 1B**). Sleep in each RNAi genotype was normalized against an RNAi control for the given experiment to control for inter-experimental differences. While this screen revealed numerous RNAi lines with an impact on sleep parameters, we focused on a single gene, *beat-Ia*, for which two independent RNAi increased sleep duration greater than 2 standard deviations of the mean. *beat-Ia* is a predicted homolog of *CADM2*, a gene within the same TAD as the non-coding variant, rs1601440, which has been reported to be associated with both excessive daytime sleepiness and daytime napping in humans ^26^ (**Fig. 1C**).

### Neuronal knockdown of *beat-Ia* increases sleep without affecting locomotor activity

CADM2 and *beat-Ia* are members of the immunoglobulin containing superfamily, functioning as synaptic adhesion molecules with known roles in neural circuit wiring^31–35^. We first replicated our findings from the screen with a larger sample size, yielding a consistent increase in sleep duration during both day and night in *beat-Ia* knockdown flies (**Fig. 2**). We also validated that both *beat-Ia* RNAi effectively knockdown mRNA expression in brain (**Fig. S1**). Excess sleep in *beat-Ia* knockdown flies was driven by an increase in sleep bout length, suggestive of more consolidated sleep (**Fig. 2F, 2I**). These initial experiments were conducted in *Drosophila* activity monitors (DAMs) utilizing a single infrared beam to detect periods of sleep, so we next assessed sleep phenotypes with *beat-Ia* RNAi using a higher spatial resolution multi-beam sleep assay. We found again that neuronal knockdown of *beat-Ia* resulted in increased sleep duration during the day and night (**Supplemental Fig. 1B-C, 1F**). Importantly, in both single- and multi-beam analyses, we detected no impact of beat-Ia knockdown on overall wake activity (**Fig. 2B, Supplemental Fig. 1E, 1H**), confirming that increased sleep is not a confound related to altered locomotor function.

**Fig. 2.**
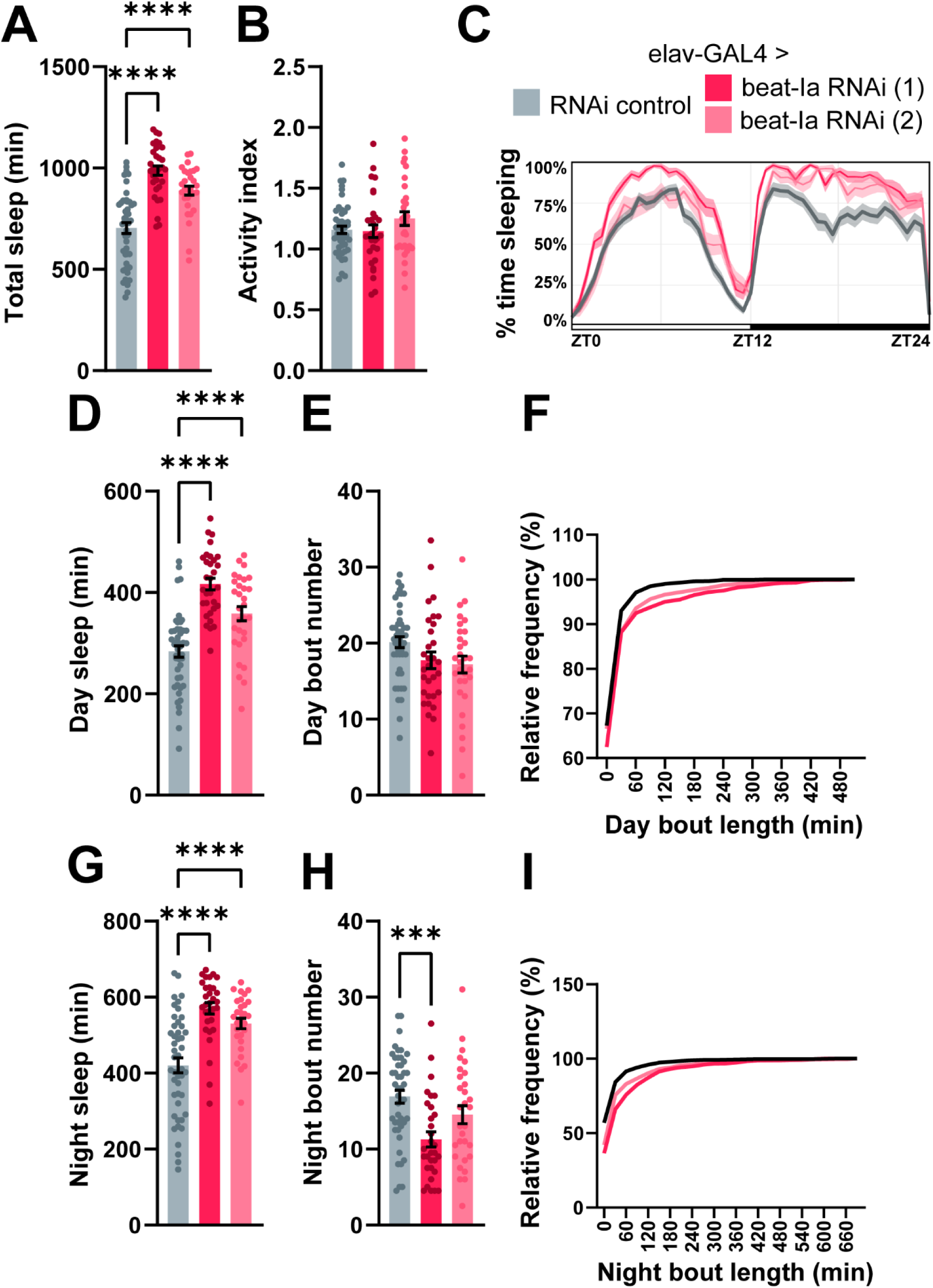
Pan-neuronal knockdown of *beat-Ia* increases sleep duration without disrupting locomotor activity. Sleep measures for flies expressing either an RNAi control or RNAi against *beat-Ia* in all neurons. (A) Total sleep duration across 24 hours. (B) Activity index, which represents the number of infrared beam breaks per minute of waking activity. (C) Sleep trace depicts sleep amount (%) in rolling 30 min bins across day (ZT0-12, light) and night (ZT12-24, dark). (D) Average daytime sleep duration. (E) Average number of daytime sleep bouts. (F) Cumulative frequency plot of the relative frequency of daytime sleep bouts of increasing duration. (G) Average nighttime sleep duration. (H) Average number of nighttime sleep bouts. (I) Cumulative frequency plot of the relative frequency of nighttime sleep bouts of increasing duration. One-way ANOVA with Dunnett’s multiple comparisons tests comparing all experimental genotypes to control. n, from left to right: 46, 31, 30. For this and following figures, * p ≤ 0.05, ** p ≤ 0.01, *** p ≤ 0.001, **** p ≤ 0.0001, and bar plots display mean +/ SEM.

Other synaptic cell adhesion molecules have been implicated in sleep regulation^36^, and *beat-Ia* is one member of the beaten path family of synaptic adhesion factors ^37^. To determine whether neuronal knockdown of other beat family members likewise affects sleep, we re-examined candidates from our initial human-informed screen. Indeed, we observed sleep phenotypes with knockdown of other beat isoforms, including increased sleep from *beat-Ic* and *beat-VII* knockdown; and decreased sleep from *beat-Ib*, *beat-IIIc*, and *beat-Vb* knockdown (**Supplemental Fig. 2**). For mechanistic studies we focused on *beat-Ia* as this phenotype was most robust and reproducible. Moreover, knockdown of *beat-Ia*’s putative receptor *sidestep* ^38,39^ (**Fig. 3**) or the transcription factor *SoxN* ^40^ (**Supplemental Fig. 3**) known to regulate *beat-Ia* expression recapitulates the hypersomnia phenotype, strongly implicating the *beat-Ia* signaling pathway in sleep.

**Fig 3.**
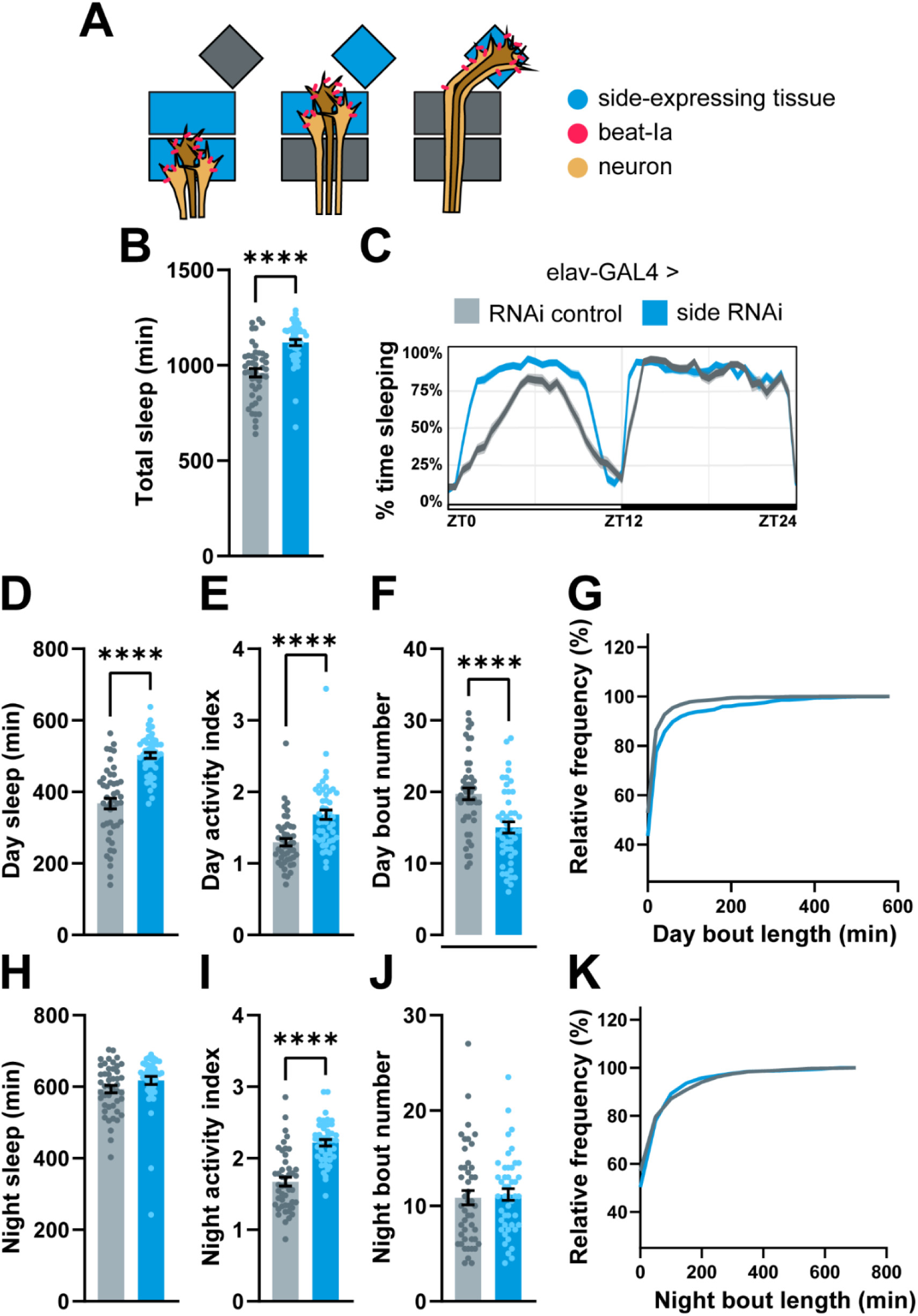
Pan-neuronal knockdown of *beat-Ia*’s receptor, side, produces a long sleep phenotype. (A) Schematic of the beat-side signaling pathway, illustrating how *beat-Ia*-expressing neurons follow a path of tissue (glia, muscle, neural, and epithelial) marked by its receptor *side*. Adapted from Siebert et al, 2009. (B) Total sleep duration. (C) Sleep trace. (D-G) Day sleep measures. (H-K) Night sleep measures. Welch’s t test. n=46 for each genotype.

### *beat-Ia* knockdown recapitulates features of human hypersomnia

Human hypersomnia is characterized by excessive sleepiness, beyond simply an increase in sleep duration. We next asked whether *beat-Ia* knockdown animals recapitulate other hypersomnia symptoms such as deeper sleep, difficulty waking from sleep, and reduced sleep onset latency. First, we analyzed our existing sleep data for additional relevant metrics. Using conditional probabilities applied to locomotor measures ^41^, we determined the probability of transitioning from wake to sleep, P(doze), or from sleep to wake, P(wake), with *beat-Ia* knockdown (**Supplemental Fig. 4**). This approach revealed a decrease in P(wake) with no change in P(doze) (**Supplemental Fig. 4B**), suggesting a reduction in probability of leaving the sleep state in *beat-Ia* knockdown flies. Notably, the elevation in sleep duration in these animals might occlude the ability to detect an increased probability of entering a sleep state, since the flies are much less commonly awake to begin with. Recent work in flies has also demonstrated that long sleep bouts reflect deeper sleep stages ^42^, so we next assessed sleep comprised of long bouts (>60 minutes) with *beat-Ia* knockdown (**Supplemental Fig. 5**). With analysis restricted to long sleep bouts, *beat-Ia* knockdown animals also exhibited longer sleep duration than control flies. Moreover, *beat-Ia* knockdown animals spend a greater percentage of time in long sleep bouts than controls, suggesting more time spent in deeper sleep (**Supplemental Fig. 5I-K**).

Next, we empirically tested other sleep propensity measures. We attempted to deprive animals of sleep using a well-established mechanical sleep deprivation protocol ^43^. This protocol dramatically reduced sleep in control animals but the effect was diminished in *beat-Ia* knockdown animals, indicative of increased resistance to sleep loss (**Fig. 4A**). We then directly assessed arousal threshold as a measure of sleep depth and found that *beat-Ia* animals were less likely to be awoken by a stimulus during sleep in comparison to controls (**Fig. 4B**). Given that individuals with hypersomnia exhibit difficulty staying awake and rapid transition to sleep, we also assessed whether *beat-Ia* knockdown animals similarly fall asleep more quickly than controls. We measured latency to sleep after animals first wake in the morning after lights turn on (ZT0). Typically, control flies first return to sleep ∼60 minutes after morning awakening; in contrast *beat-Ia* RNAi flies exhibited a first sleep bout in half that time (**Fig. 4C**). Finally, circadian dysregulation has been proposed as a mechanism of human IH. However, we observed no change to any measure of locomotor rhythms in *beat-Ia* flies under constant darkness (**Fig. 4D-E**), indicating that circadian function is unaffected and does not account for the sleep phenotypes. Taken together, loss of *beat-Ia* is associated with not only excessive sleep duration, but also numerous other sleep propensity phenotypes similar to those observed in human hypersomnia.

**Fig 4.**
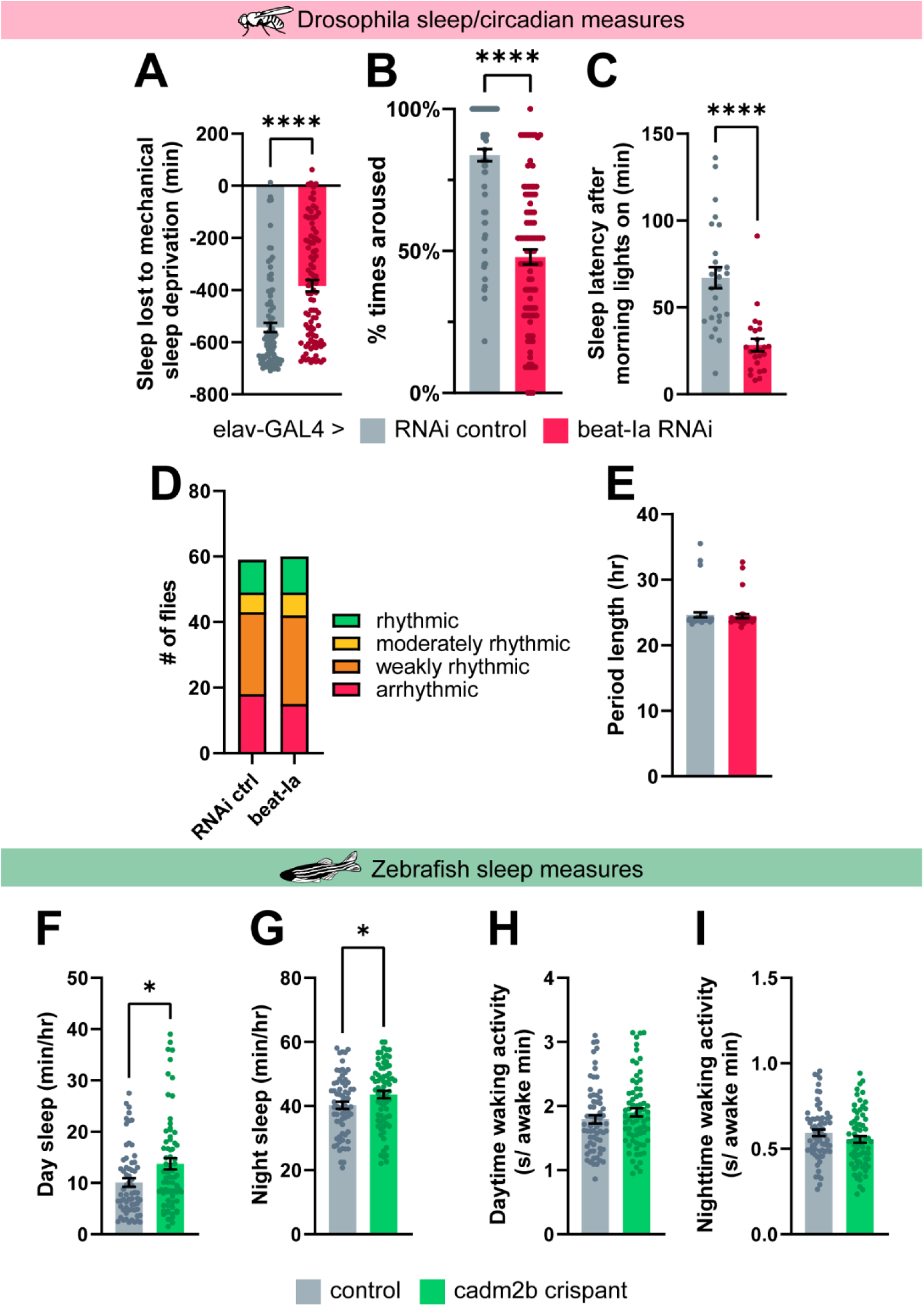
Increased sleep duration and measures of sleep propensity with loss of *beat-Ia* and *cadm2b* in flies and fish. (A) Sleep loss during 12h of mechanical sleep deprivation (measured as baseline nighttime sleep – nighttime sleep with deprivation). Welch’s t test. n, from left to right = 91, 95. (B) Arousal threshold in *beat-Ia* RNAi vs control (% of flies woken from sleep with a uniform stimulus). Welch’s t test. n, from left to right = 95, 96. (C) Sleep latency in *beat-Ia* RNAi vs control flies, following awakening at lights-on. Welch’s t test. n, from left to right = 26, 24. Analysis of rest:activity rhythms in *beat-Ia* RNAi vs controls: (D) Number of animals categorized as arrhythmic to rhythmic under constant dark conditions. n, from left to right = 59, 60. (E) Length of circadian periods in animals under constant conditions. Arrhythmic animals were excluded from analysis. n, from left to right = 42, 45. (F-I) Sleep measures for control and *cadm2b* crispant zebrafish. Mann-Whitney test. n, from left to right = 65, 71.

### *beat-Ia*’s ortholog, CADM2, regulates sleep in zebrafish

*beat-Ia* was identified as an ortholog of vertebrate *CADM2*. While *beat*s and *CADM2* are not structurally similar, they are functionally similar: both are synaptic adhesion factors of the immunoglobulin superfamily, and both are critical for axon pathfinding ^31,32,34,37,44–46^. We next tested whether vertebrate CADM2 has a role in sleep regulation by disrupting its expression in zebrafish (*Danio rerio*), a model of vertebrate sleep ^47–50^. Using CRISPR/Cas9 gene editing in F0 larvae followed by automated video sleep tracking ^8,51^, we generated exonic mutations of critical regulatory domains, predicted to produce loss-of-function through frameshifts in *cadm2b*, the most conserved transcript (**Supplemental Fig. 6**). We then screened larvae for sleep phenotypes at five days post fertilization (dpf), comparing *cadm2b* knockout zebrafish to control zebrafish (scrambled gRNA-injected) larvae. *cadm2b* knockout increased daytime and nighttime sleep, without an effect on activity, similar to sleep with *beat-Ia* knockdown in *Drosophila* (**Fig. 4F-I**) These findings support the hypothesis that CADM2 regulates vertebrate sleep.

### *beat-Ia* acts during development to regulate adult sleep

Previous studies of *beat-Ia* have focused on its role patterning the larval neuromuscular junction ^31^. In development, beat-expressing motor neurons follow a path of muscle, glial, and neuronal tissue marked by its receptor *side* ^38,39,52^. In *beat-Ia* or *side* null mutants, motor axons fail to defasciculate from the axonal bundle and do not innervate their muscular targets ^31^. *beat-Ia* null mutants are pre-adult lethal, preventing mutant analysis of adult sleep and suggesting that knockdown, as opposed to complete loss of expression, uncovers previously unknown adult roles. Although unlikely given the lack of locomotor phenotypes with pan-neuronal *beat-Ia* knockdown (see **Fig. 2, Supplemental Fig. 1**), we next asked whether the adult sleep phenotype might be related to the canonical role for this protein in motor neurons. Expression of *beat-Ia* RNAi under control of any of three well-characterized motor neuron GAL4 drivers had no effect on sleep (**Fig. 5A-B**), further supporting the hypothesis that *beat-Ia* acts elsewhere to regulate adult sleep.

**Fig 5.**
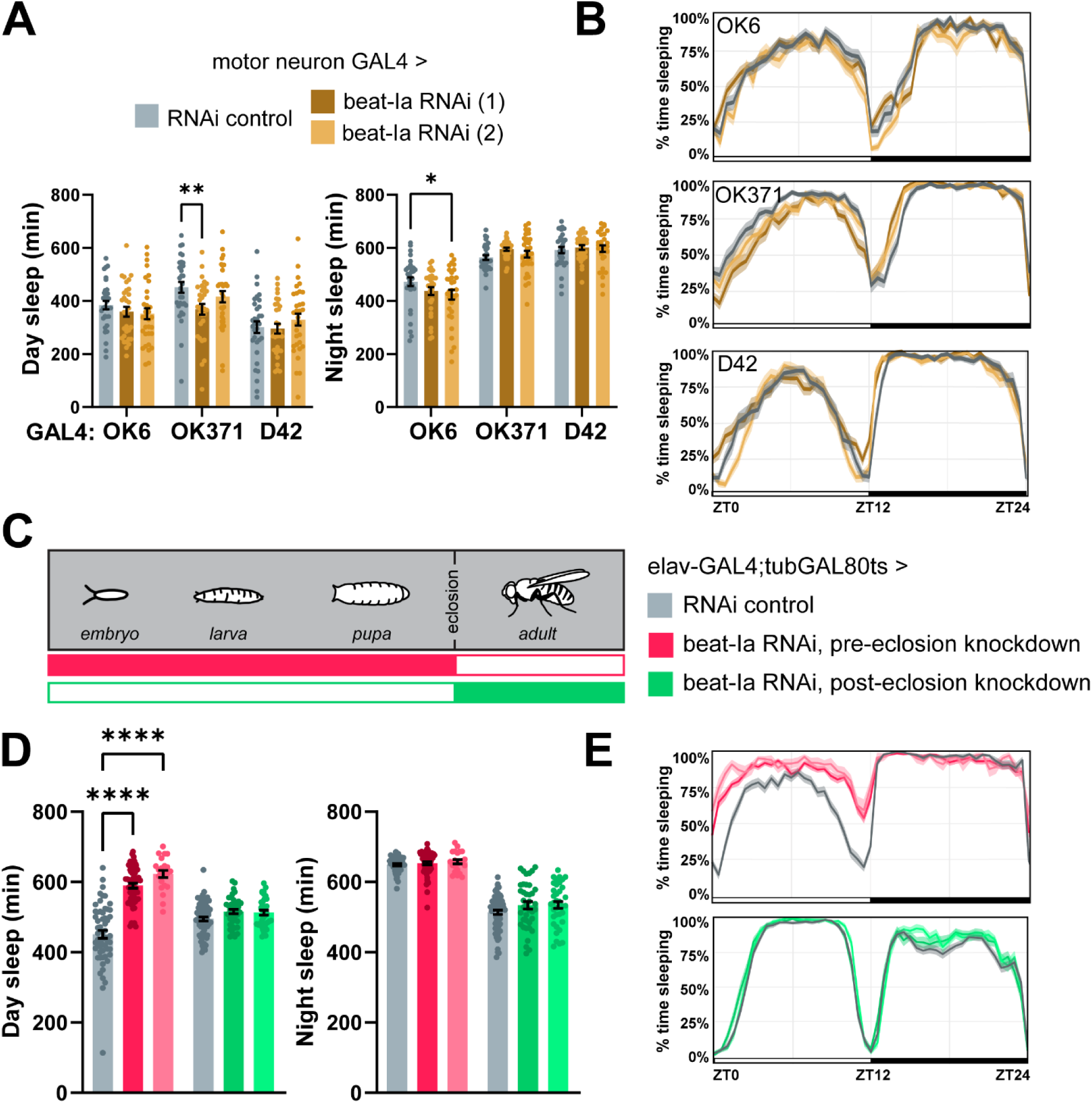
*beat-Ia* acts in development to regulate adult sleep. Day and night sleep duration (A) and sleep traces (B) from flies expressing *beat-Ia* RNAi under motor neuron drivers. Two-way ANOVA with Dunnett’s multiple comparisons tests comparing experimental genotypes to control. n, from left to right = 32, 31, 31, 32, 31, 28, 32, 31, 30. (C) Schematic of temporal restriction of *beat-Ia* knockdown using a temperature sensitive ubiquitously expressed GAL4 repressor (tubGAL80ts). Day and night sleep duration (D) and sleep traces (E) from flies in which neuronal *beat-Ia* is knocked down in development or adulthood. Two-way ANOVA with Šídák’s multiple comparisons tests comparing experimental genotypes to control. n, from left to right =52, 33, 10, 57, 38, 32.

We then tested whether *beat-Ia* is acting in development or adulthood to control adult sleep. We used a ubiquitously expressed temperature sensitive GAL4 repressor (tubGAL80ts) ^53^ to restrict *beat-Ia* knockdown to pre- or post-eclosion. First, we limited knockdown of *beat-Ia* to the pre-eclosion developmental period and then assessed sleep in adulthood. Developmental loss of *beat-Ia* was sufficient to cause the excessive sleep phenotype (**Fig. 5D-E**). In contrast, limiting *beat-Ia* knockdown to adulthood had no effect on sleep (**Fig. 5D-E**). Together, these findings demonstrate *beat-Ia* acts in a developmental capacity to cause excess sleepiness later in life.

### *beat-Ia* functions in neuropeptide F (NPF) cells to regulate sleep

Our data raise the possibility of a novel role for *beat-Ia* signaling outside of the neuromuscular junction. Given that *beat-Ia* is necessary in development for normal adult sleep, we hypothesized that it is involved in patterning adult sleep circuits, presumably in the brain. First, we showed with qPCR that *beat-Ia* is expressed in the brain (**Supplemental Fig. 1A**). To define the neuronal subpopulations in which *beat-Ia* acts to regulate sleep, we conducted a targeted spatial knockdown screen, driving *beat-Ia* RNAi in cells known to have a role in sleep or circadian behavior. We expressed *beat-Ia* RNAi in relevant neurotransmitter and neuropeptide populations, clock cells, and other sleep-implicated neuronal populations with GAL4s that express during the developmental time period (larval/pupal) in which *beat-Ia* is acting (**Fig. 6A, Supplemental Fig. 7**). Of the lines screened, we observed increased sleep when driving *beat-Ia* RNAi in cholinergic cells using ChaT-GAL4 (sleep = 1132 ± 19.02 min) compared to driving an RNAi control (sleep = 998.9 ± 32.52 min). We also observed a trend toward increased sleep when driving *beat-Ia* RNAi in neuropeptide F (NPF) cells using NPF-GAL4 (sleep = 1053 ± 23.73 min) compared to driving an RNAi control (sleep = 958.7 ± 39.55 min). ChaT-GAL4 expresses broadly throughout the fly nervous system, and we found expression overlap with NPF-GAL4 (**Supplemental Fig. 8**). Although we cannot rule out the possibility that *beat-Ia* signaling in other cells could contribute to adult sleep phenotypes, we focused on a role in NPF-expressing neurons given that NPF-GAL4 labels a relatively sparse population of cells with a known sleep regulatory role ^54–56^.

**Fig 6.**
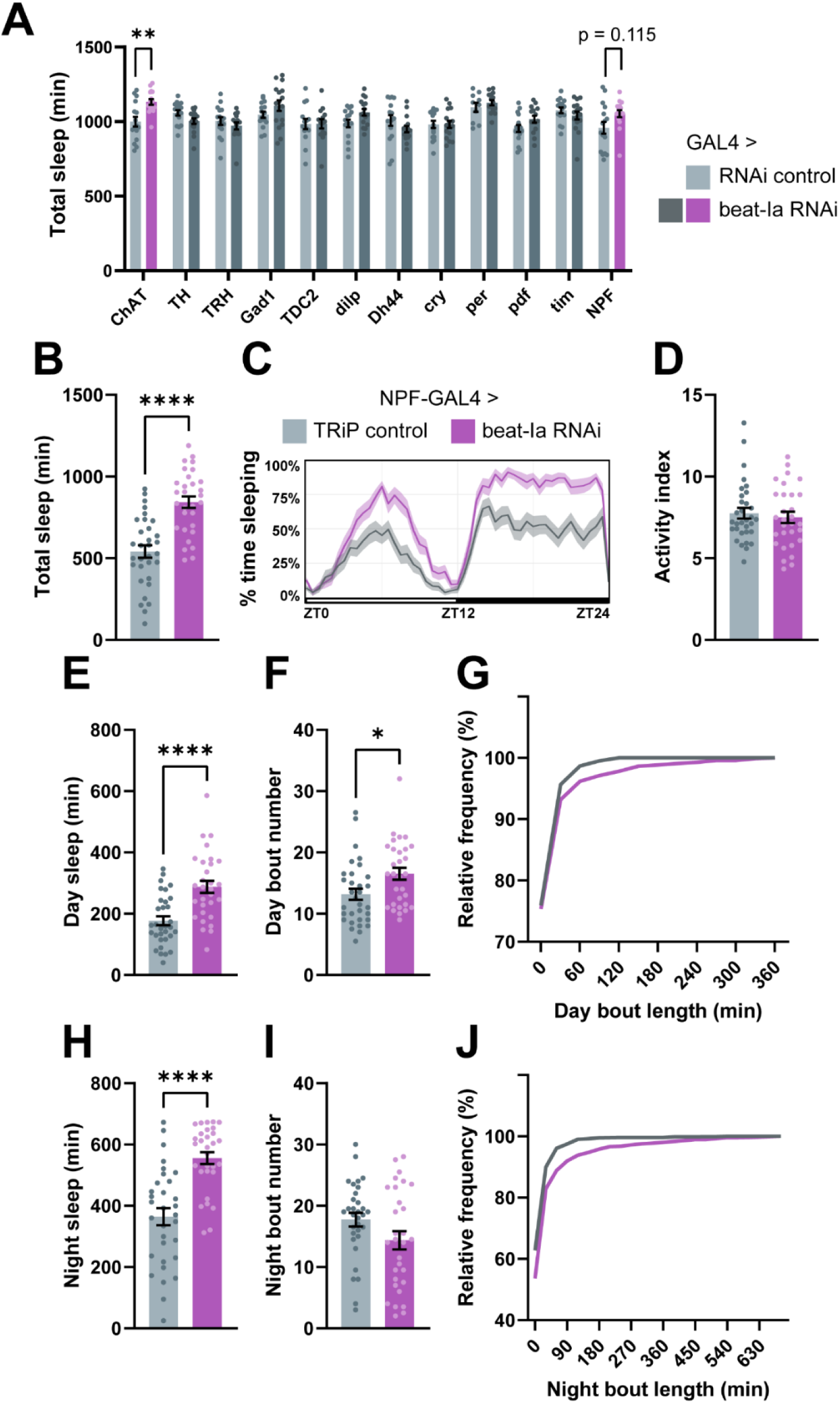
A targeted spatial knockdown screen reveals a role for *beat-Ia* in neuropeptide F (NPF) cells. (A) Total sleep duration for flies expressing *beat-Ia* RNAi or control in known sleep- and circadian-regulatory cell populations. Two-way ANOVA with Šídák’s multiple comparisons tests comparing experimental genotypes to control. N = 16 except for TRH-GAL4>*beat-Ia* RNAi (15), TDC2-GAL4>RNAi control (14), TDC2-GAL4>*beat-Ia* RNAi (14), per-GAL4>RNAi control (10), per-GAL4>*beat-Ia* RNAi (15), and pdf-GAL4>*beat-Ia* RNAi (15). (B) Total sleep duration. (C) Sleep trace. (D) Total activity index. (E-G) Day sleep measures. (H-J) Night sleep measures. Unpaired t test. n, left to right = 30, 32. Cell populations targeted: ChAT = cholinergic cells, TH = dopaminergic cells, TRH = serotonergic cells, Gad1 = GABAergic cells, TDC2 = octopaminergic cells, dilp = insulin-like peptide-secreting cells, Dh44 = diuretic hormone 44-producing cells, cry, per, pdf, tim = clock cells, NPF = neuropeptide F-producing cells.

Following the initial candidate screen, we replicated the NPF results using higher spatial resolution sleep monitoring, which confirmed a strong excess sleep phenotype (**Fig. 6B-J**). As in pan-neuronal *beat-Ia* RNAi flies, knockdown in NPF cells was not only associated with increased day and night sleep (**Fig. 6E, H**), but also prolonged sleep bouts (**Fig. 6G, J**). Importantly, locomotor activity was unaffected (**Fig. 6D**). Analysis of long sleep bouts, a proxy for deeper sleep ^42^, revealed a dramatic increase in this type of sleep during day and night, as well as increased probability of even lengthier sleep bouts than controls (**Supplemental Fig 9**). Thus, *beat-Ia* RNAi expression in NPF cells recapitulates sleep features observed with pan-neuronal knockdown.

NPF cells have a known role in sleep and feeding, among other behaviors in the fly. Previous work has shown that activation of NPF cells can promote wakefulness and feeding, and that these functions are dissociable: specific NPF neuronal subpopulations promote wakefulness without affecting feeding, and vice versa ^55^. We hypothesized that the increase in sleep in NPF>*beat-Ia* RNAi animals is due to a loss of a wakefulness-promoting cue. Alternatively, NPF>beat RNAi animals could experience altered feeding behavior or nutrient storage, leading to disrupted energy homeostasis and an indirect effect on sleep. However, NPF>*beat-Ia* RNAi flies showed no defects in feeding or nutrient storage as measured by the Capillary Feeding Assay and a starvation resistance assay ^57^ (**Supplemental Fig. 10**), supporting a specific developmental role for *beat-Ia* in NPF arousal circuitry.

### *beat-Ia* coordinates NPF neuronal synaptic development in the suboesophageal zone (SEZ)

To determine how loss of *beat-Ia* affects NPF wiring, we first used NPF-GAL4 to co-express membrane-bound GFP (UAS-CD8::GFP) with *beat-Ia* RNAi or an RNAi control. Surprisingly, we observed no changes in NPF cell body number or neurite morphology/location (**Fig. 7A**). NPF cells have dense dendritic and axonal projections, precluding assessment of more subtle morphological effects using this approach. Given that *beat-Ia*’s known neuronal function is presynaptic, we next labeled presynaptic sites using EGFP tagged Synaptotagmin (Syt-EGFP). While co-expression of *beat-Ia* RNAi had no detectable impact on most regions where NPF cells send projections, we noted a dramatic loss of synaptic density in the suboesophageal zone (SEZ) of the fly brain (**Fig. 7B-C**). Closer examination of axonal projections confirmed that NPF axons reach the SEZ and appear morphologically normal (**Fig. 7A**, inset), suggesting a defect in synaptogenesis or synaptic maintenance in NPF projections to the SEZ with loss of *beat-Ia*. We therefore assessed when NPF neurons normally innervate the SEZ. Imaging of NPF>Syt-EGFP brains across pupal development, we found elaboration of NPF neuronal presynaptic sites in the SEZ occurs during mid- to late-pupal development (**Supplemental Fig. 11**). Since pan-neuronal knockdown of *beat-Ia*’s receptor *side* also results in excess sleepiness in adulthood (see **Fig. 3**), we wondered whether *side* might be involved in the *beat-Ia*-dependent SEZ phenotype of NPF projections. Strikingly, we found that *side* is specifically expressed in ∼6-8 cells in the SEZ region during the mid-pupal stage (**Fig. 7C**); this expression dissipates and is absent in adulthood. These findings support a model in which transient expression of *side* in the SEZ coordinates proper NPF synaptic connectivity with downstream targets during development, essential for normal arousal in adulthood.

**Fig 7.**
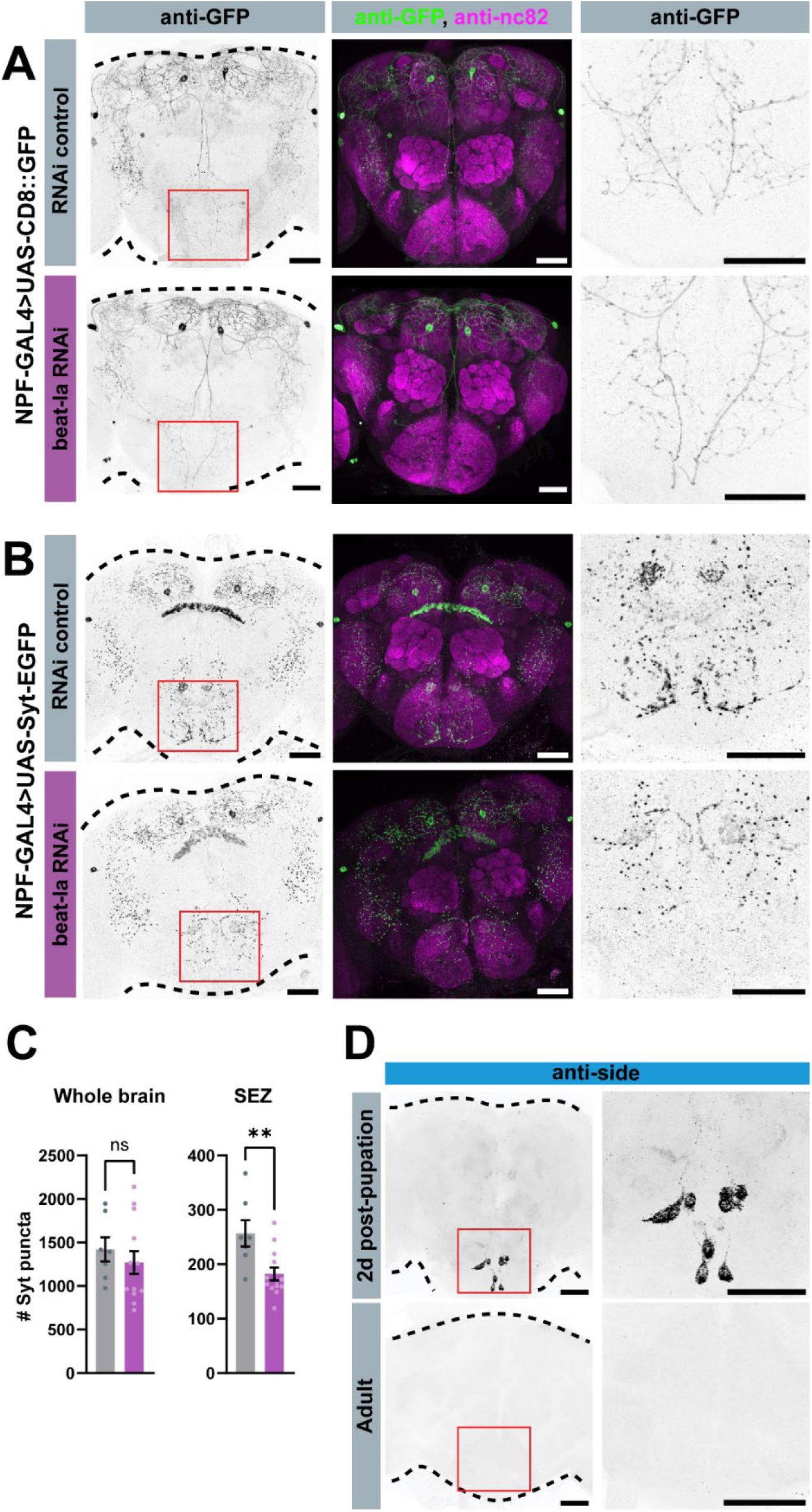
*beat-Ia* knockdown in NPF cells disrupts synaptic development in the suboesophageal zone (SEZ). (A-B) Representative images from adult brains of flies expressing membrane reporter CD8::GFP (A) or presynaptic reporter Syt-EGFP (B) under control of NPF-GAL4 with or without beat-Ia RNAi. All images are maximum projections except for SEZ insets of CD8::GFP brains, which are projections of z-stacks cropped to only show SEZ signal. In SEZ insets of Syt-EGFP images (B), *beat-Ia* RNAi animals show a marked reduction in Syt-EGFP signal. (C) Quantification of Syt-EGFP-tagged puncta across entire brain and in SEZ. Unpaired t-test. n, left to right = 7, 13. (D) Anti-side staining shows that side expression is temporally limited to pupal stages and spatially restricted to the SEZ. Scale bar = 50µM.

Finally, given that *beat-Ia* knockdown in NPF cells impairs synapse development and adult wakefulness, we examined whether *beat-Ia* overexpression might yield the opposite phenotypes. We found expression of UAS-*beat-Ia* along with Syt-EGFP in NPF neurons resulted in a dramatic increase in presynaptic sites across many areas of the brain, including SEZ (**Fig. 8A-B**). Moreover, *beat-Ia* overexpression flies exhibited a decrease in total and daytime sleep (**Fig. 8C-E**), providing further evidence that *beat-Ia*-directed NPF synaptic development tunes sleep-wake balance in adulthood.

**Fig 8.**
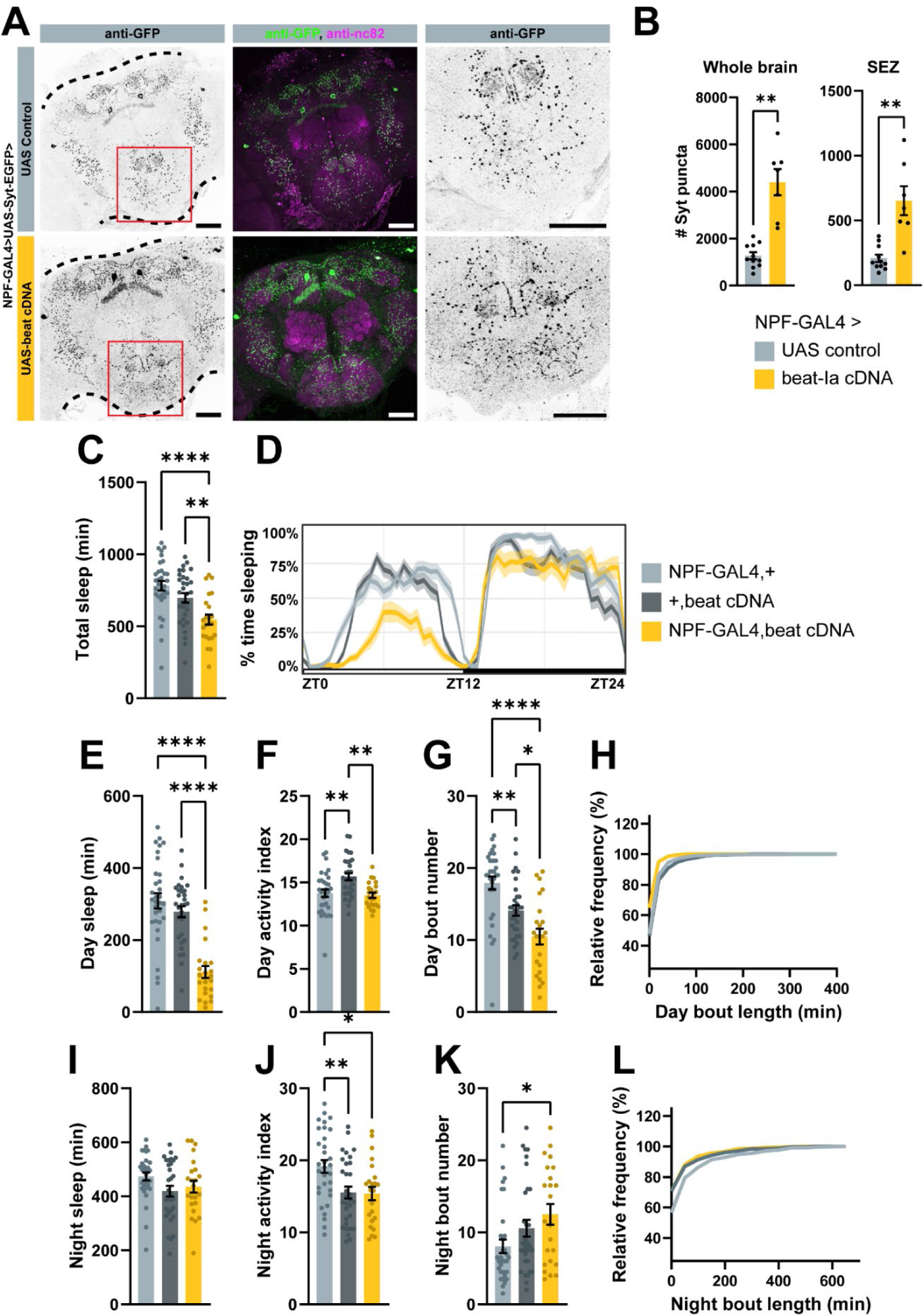
*beat-Ia* overexpression in NPF cells increases synapse number and reduces sleep. (A) Representative images from flies expressing presynaptic reporter Syt-EGFP and *beat-Ia* cDNA or a UAS control using the NPF-GAL4 driver. Scale bar = 50µM. (B) Quantification of Syt-EGFP-tagged puncta across entire brain and in SEZ. Welch’s t test. n, left to right = 11, 7. (C) Total sleep duration. (D) Sleep trace. (E-H) Day sleep measures. (I-L) Night sleep measures. One-way ANOVA with Tukey’s multiple comparisons tests to compare experimental genotype to controls. n, from left to right = 32, 31, 23.

### Identification of putative downstream sleep-regulatory cells in the SEZ

If loss of NPF synaptic innervation in the SEZ causes excess sleep in *beat-Ia* knockdown animals, then presumably NPF cells are synapsing onto sleep-regulatory SEZ targets. What are the downstream neuronal targets that would have received NPF input if not for loss of *beat-Ia*? Using FlyWire, a publicly available *Drosophila* connectome dataset ^58,59^, we identified 22 candidate cells that receive input from either of the descending NPF neurons (SLP.AVLP.4 and SLP.AVLP.5, in FlyWire) and have input regions in the SEZ (“gnathal ganglion” in the FlyWire dataset) (**Fig. 9A**). Using BrainCircuits.io ^60^, we then identified sparsely expressing GAL4 driver lines with expression predicted to match at least one of these cells of interest. Of the 62 GAL4s identified, 17 drivers were matched to at least 3 cells each and altogether covered the majority of putative downstream cells (77%), and 11 were publicly available (**Supplemental Data 2**).

**Fig 9.**
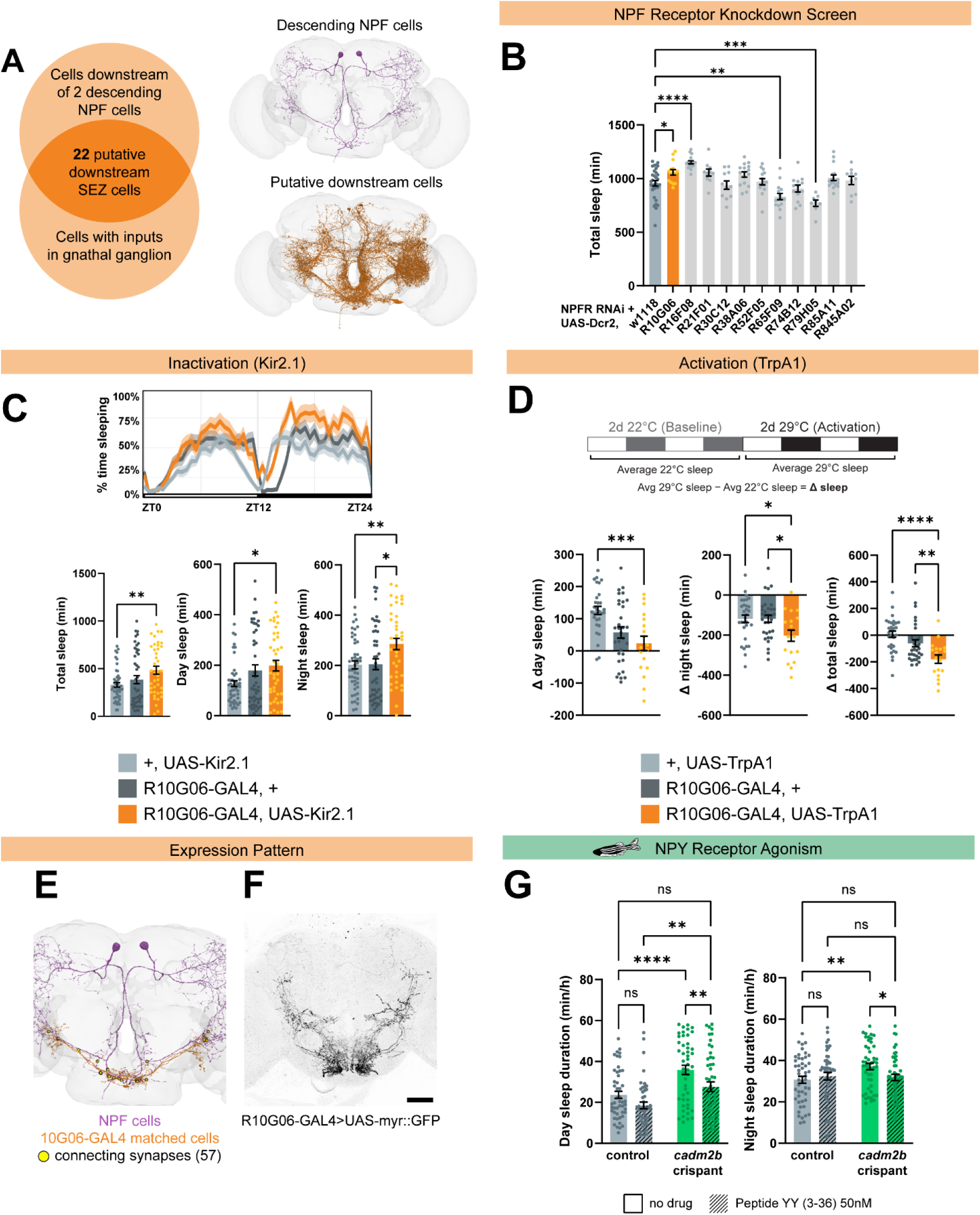
Identification of downstream SEZ cells that regulate sleep. (A) Schematic of connectome-assisted identification of putative SEZ cells downstream of NPF. (B) Initial screen of sleep with NPF receptor knockdown in SEZ cells. n, left to right = 32, 15, 16, 10, 11, 16, 15, 15, 13, 8, 16, 10. (C) Sleep with inactivation of 10G06 cells using the inwardly rectifying potassium channel Kir2.1. n, from left to right = 40, 48, 47. (D) Sleep with activation of 10G06 cells with heat-sensitive cation channel TrpA1. Change in sleep due to activation was calculated by taking the difference of each sleep measure at 29C from sleep at 22C. n, left to right = 18, 30, 32. One-way ANOVA with Dunnett’s multiple comparisons tests comparing experimental genotypes to controls. (E) EM-reconstructed traces of NPF cells and cells matched to 10G06-GAL4. Images taken from Neuroglancer. (F) Expression pattern in adult brain of 10G06-GAL4. Scale bar = 50µM. (G) Day and night sleep duration in negative control siblings and *cadm2b* crispant zebrafish with or without NPY Y2 receptor agonism by Peptide YY (3-36). Two-way ANOVA with Uncorrected Fisher’s LSD comparing within genotypes no drug to Peptide YY, within drug conditions negative control siblings to *cadm2b* crispants, and negative control siblings with no drug to *cadm2b* crispant on Peptide YY. n = 48 for all genotypes.

If *beat-Ia* knockdown in NPF cells increases sleep by reducing NPF input into the SEZ, then knocking down the NPF receptor (*NPFR*) in downstream cells should also increase sleep. We expressed *NPFR* RNAi using each of the candidate GAL4s and found excess sleep with two drivers, R10G06 and R16F08, that label 3 overlapping cells in the SEZ (**Fig. 9B**). Following this initial candidate screen, we replicated experiments and found a consistent increase in sleep with R10G06>*NPFR* RNAi (**Supplemental Fig. 13**), while the phenotype with R16F08 was only significantly different compared to one parental control (**Supplemental Fig. 14**). *NPFR* knockdown in R10G06 cells also increased time spent in long sleep bouts (**Supplemental Fig. 15**), demonstrating that reduced NPF signaling onto these cells increases both sleep duration and sleep depth. The NPF circuit normally promotes or maintains arousal, so manipulating these cells downstream of NPF input would be predicted to modulate sleep. Inhibition of cells defined by R10G06 using *UAS-Kir2.1*, an inwardly rectifying potassium channel, resulted in increased sleep (**Fig. 9C**), similar to *NPFR* knockdown or loss of NPF input via *beat-Ia* RNAi (R16F08>Kir2.1 was lethal). We then tested activation of R10G06 cells with the heat-sensitive cation channel *UAS-TrpA1* and observed a reduction in sleep (**Fig. 9D**), supporting a role in arousal. Further connectome analysis revealed that R10G06-GAL4 (and R16F08-GAL4) expresses in three specific candidate cells of the SEZ downstream of NPF neurons (GNG.590, GNG.743, and GNG.651), all of which are GABAergic and project almost exclusively to other cells within the SEZ (**Fig. 9E,F**). Together, these data support a model in which a subpopulation of ventrally-projecting NPF cells activate three GABAergic neurons in the SEZ to maintain wakefulness, and that excessive sleep can emerge from developmental miswiring of this circuit (**Supplemental Fig. 16**).

### NPY receptor agonism rescues long sleep phenotype in *cadm2b* crispant zebrafish

Our evidence in flies defines a developmental origin of adult hypersomnia but implicates a specific signaling system (NPF) whose dysfunction can be targeted. To avoid confounds related to oral administration of peptides in flies, we examined pharmacologic manipulation of NPY signaling in *cadm2b* crispant zebrafish, in which drugs can be bath applied ^61^. While *Drosophila* express only one isoform of the NPF receptor, zebrafish express five NPY receptor subtypes ^62^, including an NPY Y2 isoform that is highly homologous to both human NPY Y2 and *Drosophila* NPF receptor ^63^. We administered a selective agonist of the human NPY Y2 receptor, Peptide YY (3-36) ^64,65^, and measured sleep in *cadm2b* crispants and sibling negative control-injected fish. As expected, without Peptide YY (PYY) administration, *cadm2b* crispants slept significantly more than controls (**Fig. 9G**). Remarkably, with PYY administration, excess sleep during both day and night in *cadm2b* crispants was attenuated, reaching similar sleep duration to control fish (**Fig. 9G; Supplemental Fig. 17**). PYY exposure in controls had no significant effects on sleep, supporting specific efficacy of the drug in the setting of hypersomnia with loss of *cadm2b* (**Fig. 9G**). These findings suggest conserved impairment of NPF/NPY signaling with loss of *beat-Ia/cadm2b* and point towards NPY agonism as a novel therapeutic avenue in human hypersomnia.

## Discussion

Disorders of excessive sleepiness are poorly studied, with few mechanistic insights. Leveraging a comprehensive TAD-wise analysis of putative candidate genes related to GWAS-implicated hypersomnia loci, we describe a conserved role for synaptic adhesion molecules *beat-Ia* and *CADM2* in sleep regulation. Specifically, *beat-Ia* patterns sleep circuits during development, guiding wiring of NPF neurons to promote normal arousal in adulthood. Our results indicate that descending NPF projections to the SEZ undergo synapse elaboration onto a small population of wake-promoting GABAergic neurons, coordinated by *beat-Ia*’s receptor, *side*. With loss of *beat-Ia*, these synapses fail to develop normally, resulting in impairment of the arousal-stabilizing NPF signal and excessive sleepiness in adulthood (**Supplemental Fig. 16**). Pharmacologic evidence in zebrafish suggests similar NPY circuit impairment with loss of *cadm2b* that can be corrected with an NPY receptor agonist. Combining human genomics with experimental observations in fish and flies, our findings uncover a previously unrecognized neurodevelopmental basis for IH stemming from dysfunction of a targetable neuropeptidergic system.

*beat-Ia* knockdown in neuropeptide F (NPF) cells is sufficient to cause sleepiness in adult flies, through loss of NPF synaptic innervation in development and the resulting impaired NPF signaling in adulthood. NPF is the *Drosophila* homolog of vertebrate neuropeptide Y (NPY), and both NPF and NPY play a role in numerous behaviors, including feeding, social behavior, learning, circadian rhythmicity, and sleep. Both wake- and sleep-promoting roles have been described for NPF/NPY, depending on manipulation, sex, and organism. In zebrafish, global overexpression of NPY promotes sleep and loss of NPY reduces sleep ^66^. In *Drosophila*, activation of specific NPF subpopulations promotes wakefulness ^55^, while others might be sleep-promoting ^54^. Our findings, combined with previous results, support the hypothesis that such varied effects are likely due to NPF’s diverse downstream targets; *NPFR* is expressed widely across the brain ^55^, and NPFR activation could result in inhibition or excitation of either wake- or sleep-promoting cells depending on the circuit. While pharmacologic results support NPY dysfunction with loss of *cadm2b* in zebrafish, future work will aim to define how specific NPY microcircuits are affected, as parsing sleep-versus arousal-promoting roles, among other behaviors like feeding, will clarify NPY-related mechanisms of excessive sleepiness across phylogeny.

Our work also reveals that *beat-Ia* and its receptor *side* have a role in neurodevelopment in the central brain. Until now, *beaten path* signaling has only been implicated in motor neuron ^32,44^ and visual system development ^67,68^, but here we demonstrate a role for beat-side signaling in the development of central brain circuits, including but not limited to NPF cells. We focused specifically on *beat-Ia* and its receptor *side*, but knockdown of several other beat isoforms produced sleep phenotypes as well. It is possible that *beats* and *sides* act in a combinatorial manner to pattern numerous central brain circuits, as they have been predicted to act in other neural tissues ^69^.

By intersecting our experimental findings with the FlyWire *Drosophila* whole-brain connectome, we identified three GABAergic neurons in the SEZ downstream of NPF cells as part of a sleep-regulatory circuit. Knockdown of *NPFR* and/or inhibition of these cells defined by the R10G06-GAL4 driver promoted sleep, and activation promoted arousal. NPF cells likely signal onto GABAergic SEZ cells, activating these neurons which then inhibit downstream, sleep-promoting targets, resulting in wakefulness (ON→ON→OFF→wake) (**Supplemental Fig. 16**). In *beat-Ia* knockdown animals, NPF cells do not signal onto SEZ GABAergic cells, thus disinhibiting downstream sleep-promoting neurons and allowing excess sleep to occur (OFF→OFF→ON→sleep) (**Supplemental Fig. 16**). Such circuit organization is broadly similar to that of GABAergic cells in the mouse lateral hypothalamus, which inhibit sleep-promoting neurons in the ventrolateral preoptic area during wakefulness and are inactive during sleep ^70^. This model supports the hypothesis that some forms of IH result from impaired stability of arousal, and pinpoints specific neuropeptide signaling systems at play.

Finally, our work highlights a role for immunoglobulin synaptic adhesion molecules such as *beat-Ia* and *CADM2* in sleep regulation. In mammals, Ig containing neural cell adhesion molecules (NCAMs), similar to *CADM2*, have been found to act in adulthood to regulate sleep and circadian function ^36,71^, but specific mechanisms have remained elusive and developmental roles have not been described.

Neurodevelopmental disorders with links to synaptic adhesion molecule alterations ^34,72^ are commonly comorbid with sleep disturbances ^73,74^, suggesting a shared etiology. Our findings demonstrate that some sleep disorders themselves, including hypersomnias, could stem from neurodevelopmental disturbances yet remain correctable in adulthood.

## Materials and Methods

### Drosophila husbandry

Unless otherwise specified, flies were raised and maintained on standard molasses food (8.0% molasses, 0.55% agar, 0.2% Tegosept, and 0.5% propionic acid) at 25°C on a 12-hour:12-hour light:dark (LD) cycle. 5-7d old female flies were used in all experiments unless otherwise noted.

### Drosophila strains

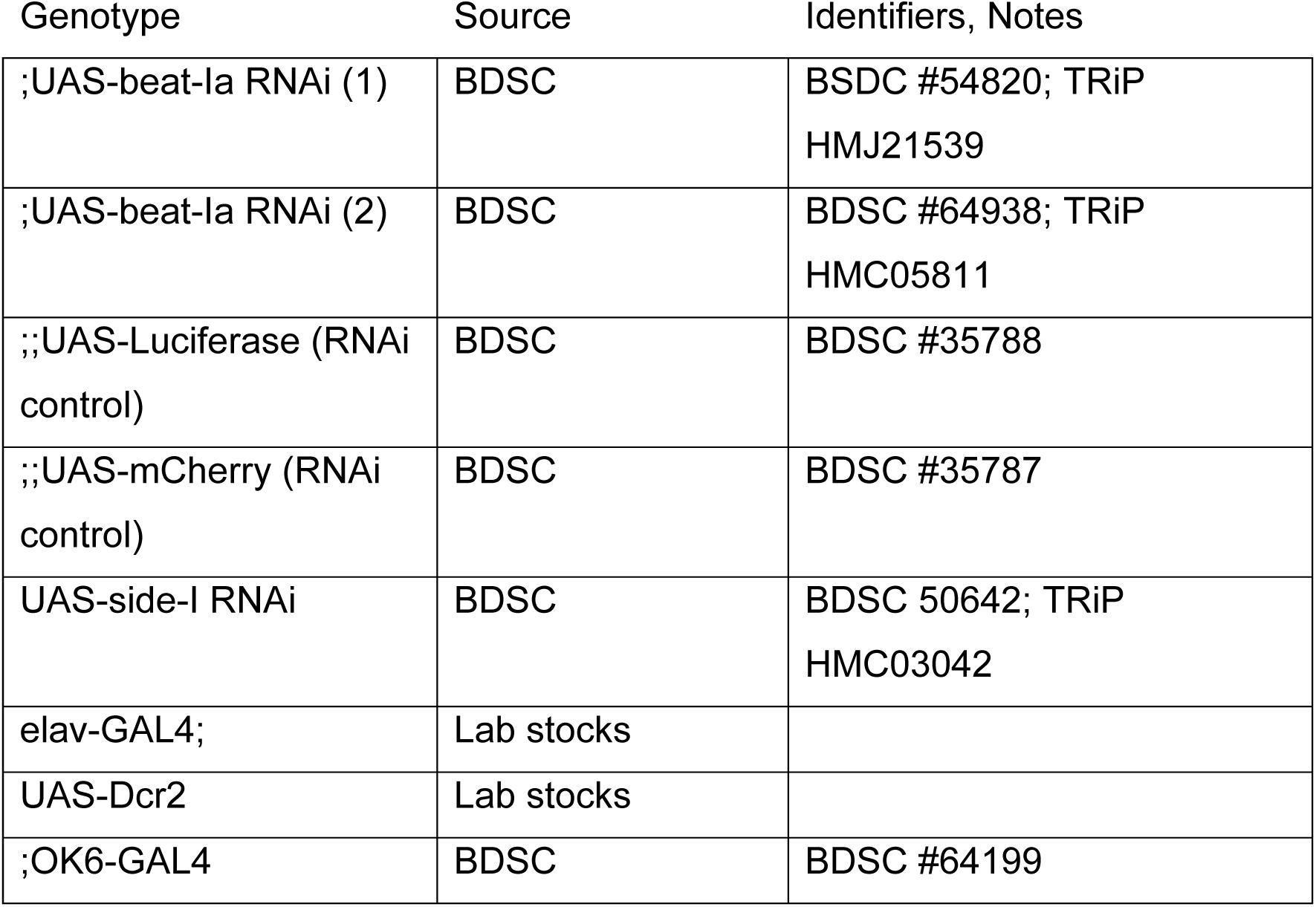
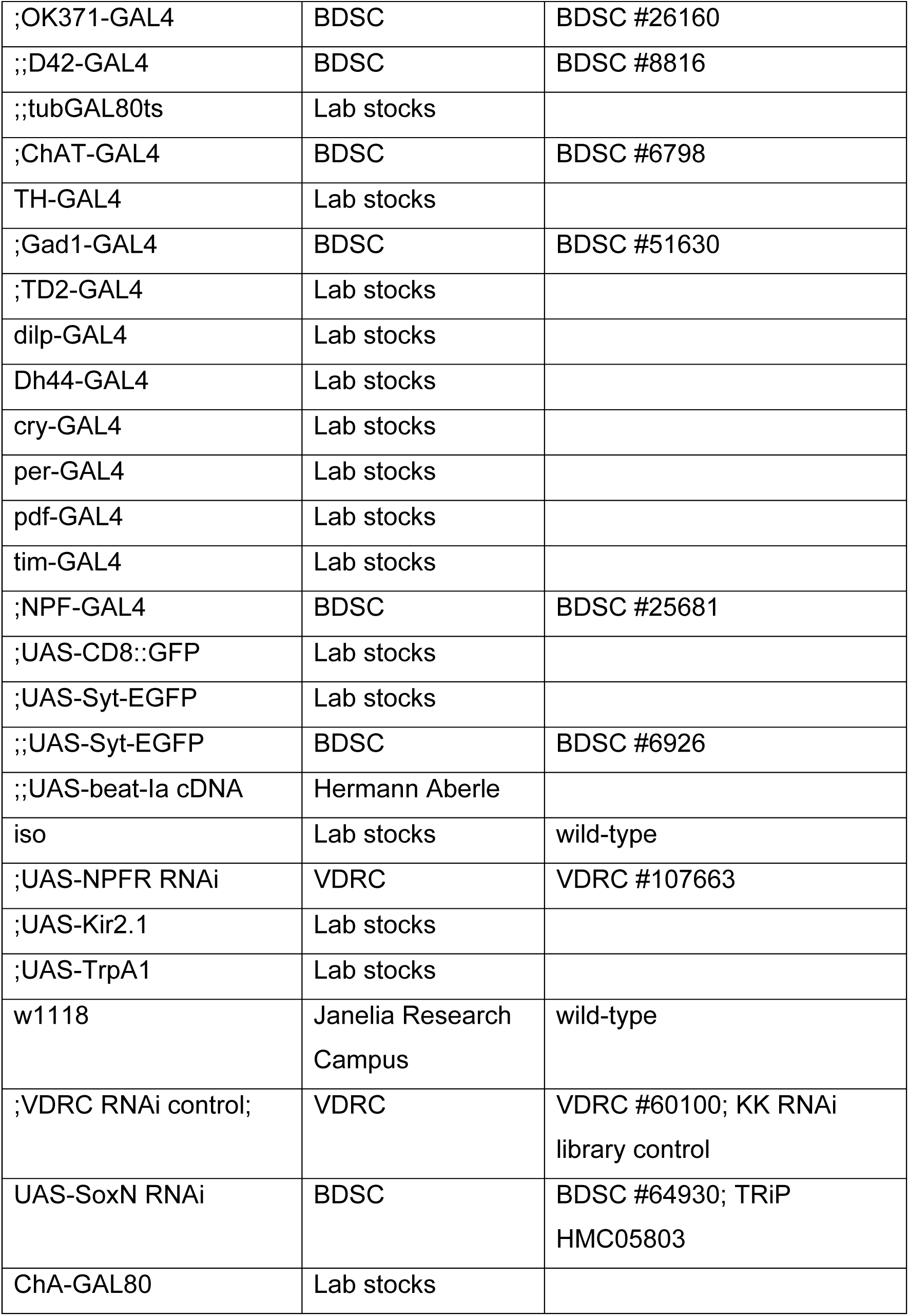

### Gene identification, TAD analysis

TAD boundaries were downloaded from http://3dgenome.fsm.northwestern.edu/^75^ for hg19 H1-NPCs^22^ to determine the extent of the TAD encompassing rs1601440. Locus zoom plots were created using the Sleep Disorders Knowledge Portal (https://sleep.hugeamp.org/) with the LD reference SNP set at rs1601440 for against the EUR reference panel.

### Sleep assays

Adult female flies were collected 2-3d after eclosion and aged to 5-7d in group housing at 25°C on a 12-hour:12-hour light:dark schedule, unless otherwise noted. Female flies were housed with males to ensure mated status upon loading. Flies were then anesthetized on CO_2_ pads (Genesee Scientific, #59-114) and loaded into individual glass tubes supplied with 5% sucrose, 2% agar food. Tubes were then loaded into Drosophila Activity Monitoring systems (Trikinetics) and flies were monitored for at least 2d. Single beam (DAM2) systems were used for Figures 1-5, 6A, 2s2, 3s1, 4s1, 4s2, and 6s1, and multibeam (DAM5H) systems were used for figures 6B-J, 8, 9, 2s1, 6s4, 8s1, 9s1, 9s2, and 9s3. Activity was measured in 1m bins and sleep was defined as 5m of inactivity. Activity indices were measured as number of beam breaks per 1m of waking activity. Data was analyzed using Rethomics.

### Sleep deprivation

Flies received a 2s stimulation randomly within every 20s window from ZT12-ZT24 using a mechanical vortexer (Trikinetics). “Sleep lost” was measured as the difference between minutes of sleep during mechanical deprivation and minutes of baseline nighttime sleep.

### Arousal threshold

Flies received a 0.5s stimulation once per hour from ZT13-ZT23, totaling 11 timepoints per animal. Flies were defined “aroused” if they were in a >5m immobility bout prior to stimulation and made a beam break within 2m after stimulation, and “not aroused” if they were in a >5m immobility bout that was not broken by stimulation. “Arousability” was measured per fly as # times aroused/# of times not aroused.

### Sleep latency

Adult female flies were loaded into the single beam DAM system as previously described. Sleep latency was measured by identifying animals that were woken by the light transition at ZT0 and measuring time lapsed until their next sleep bout, averaging sleep latency across two days. Animals that were awake at the time of the light transition or who were not woken were excluded.

### Circadian rhythmicity analysis

Flies were loaded into the DAM system 5-7d after eclosion and entrained to a 12-hour:12-hour LD cycle for 4d before transferred to constant darkness (DD). Locomotor activity was measured for 11d in DD. Locomotor rhythmicity was analyzed by Fast Fourier transform (FFT) using ClockLab software (Actimetrics, Wilmette, IL), and the maximum FFT amplitude was calculated to determine rhythmicity: rhythmic (FFT ≥ 0.05), moderately rhythmic (0.05 > FFT ≥ 0.03), weakly rhythmic (0.03 > FFT ≥ 0.01), or arrhythmic (<0.01). Period length was calculated only from flies that were not categorized as arrhythmic.

### Temporal mapping experiments

Using elav-GAL4;;tubGAL80ts to drive RNAi, flies were held at 18°C to permit GAL80 expression and block RNAi expression or were held at 29°C to denature GAL80 and allow gene knockdown. The following temperature shift schedules were used to restrict knockdown to developmental stages:

For pre-eclosion knockdown, parental crosses were raised at 29°C and immediately upon eclosion progeny were collected and moved to 18°C. Progeny were aged to 5-7d and sleep assays were conducted at 18°C.

For post-eclosion knockdown, parental crosses were raised at 18°C and immediately upon eclosion progeny were collected and moved to 29°C. Progeny were aged to 5-7d and sleep assays were conducted at 29°C.

### Generation of Zebrafish mutant

Zebrafish experiments were performed in accordance with the University of Pennsylvania Institutional Animal Care and Use Committee guidelines.

Wild-type (AB/TL) embryos were collected in embryonic growth media (E3 medium; 5mM NaCl, 0.17 mM KCl, 0.33 mM CaCl2, 0.33 mM MgSO4) in the morning shortly after lights-on. Pre-formed ribonuclear protein complexes containing the guide RNAs (gRNA) and Cas9 enzyme were injected at the single-cell stage. Negative control sequences are: 5’-CGTTAATCGCGTATAATACG-3’ and 5’-CATATTGCGCGTATAGTCGC-3’. For *cadm2b*, two gRNAs were designed using the online tool Crispor (http://crispor.tefor.net/) with the reference genome set to “NCBI GRCz11” and the protospacer adjacent motif (PAM) set to “20bp-NGG-Sp Cas9, SpCas9-HF, eSpCas9 1.1.” gRNAs were prioritized by specificity score (>95%) with 0 predicted off-targets with up to 3 mismatches. The zebrafish sequence was obtained using Ensembl (https://useast.ensembl.org/) with GRCz11 as the reference genome. Sequence was aligned to the human amino acid sequence using MARRVEL (http://marrvel.org/) to identify the coding region with highest conservation, and each gRNA was designed targeting conserved exonic regions. gRNA 1 was designed to target exon 3 of transcript 201 with the guide sequence 5’-TTGAACTGGTCCGAGCCTCG-PAM-*TGG*-3’; Left primer 5’-TCAGCATTGAGGGACAACCG-3’; Right primer 5’-ATGACTGGGAGAGAACGCAC-3’; Headloop sense primer 5’-CGAGGCTCGGACCAGTTCAATCAGCATTGAGGGACAACCG-3’. gRNA 2 was designed to target exon 4 of transcript 201 with the guide sequence (reverse strand) 3’-*CCG*-PAM-ATGGTTCAGGAACGACAAGG*-*5’; Left primer 5’-TCCTGCTGCTCTGTTTGTGT-3’; right primer 5’-ACACAGATCACGCACCACTT-3’ and restriction enzyme BccI. DNA extraction was performed per the manufacturer’s protocol (Quanta bio, Beverly, MA) immediately following completion of the sleep assay (8 days post fertilization), as described previously^8,51^. Genotyping was performed on individual fish at the conclusion of each sleep assay by PCR following manufacturer’s protocols (64° for Headloop PCR using Phusion HF DNA polymerase and 60°for restriction digest using GoTaq Green Master Mix). Mutation efficiency was quantified by band intensity as the ratio of headloop primer: standard primer and cut:uncut, and fish were included in behavioral analysis provided they show >90% efficiency in at least 1 guide RNA as described previously^51^. All primers were run on a 2% agarose gel and sequence verified using Sanger sequencing to verify the target region.

### Data collection and analysis for sleep phenotyping in zebrafish

On day 5 post fertilization, CRISPR mutants and scramble-injected sibling controls were pipetted into individual wells of a 96-well plate and placed into a Zebrabox (Viewpoint Life Sciences) for automated video monitoring. Genotypes were placed into alternating rows to minimize location bias within the plate. Each zebrabox is sound-attenuating and contains circulating water held at a temperature of 28.5°C with automated lights cycling on the same 14-hour:10-hour light/dark cycle. E3 media was topped off at lights-on (9am) every day. Sleep-wake behaviors were measured through automated video-tracking, as described previously^8,51,61^.

Activity data were captured using automated video tracking (Viewpoint Life Sciences) software in quantization mode. As described previously (Chen et al., 2017), threshold for detection was set as the following: detection threshold: 20; burst: 29; freeze: 3; bin size: 60 seconds. Data were processed using custom MATLAB scripts (Lee et al., 2022) with the threshold for sleep set at 1 min with <0.5 second of activity. All animals were allowed to acclimate to the zebrabox for approximately 24 hours before beginning continuous data collection for 48 hours starting at lights-on. Two biological replicates were performed swapping row order in the plate.

### Peptide YY (3-36) administration and sleep phenotyping

Peptide YY (3-36) (PYY) was obtained from Tocris (#1618) and dissolved in water to achieve a 100µM stock solution. Fish were allowed to develop and hatch normally without manual dechorionation. On day 3 post fertilization, larvae were placed into individual wells of a 96-well plate and E3 media was filled to be flush with the top of each well (650µl). PYY was dissolved in water and 0.65µl was administered directly to the well at a concentration of 50µM to give a final drug concentration of 50nM in each well. An equal volume of water was added to negative control wells. PYY was administered at 2pm on days 3 and 4 post fertilization when E3 was topped off in each well. Fish were allowed to acclimate to the box and sleep was measured as described above except sleep-wake behaviors were only measured on day 4 to capture acute effects of drug administration. Two biological replicates were performed swapping row order in the plate.

### CAFÉ Assay

Feeding was measured using a modified version of the Capillary Feeder assay. 10 adult female flies were transferred from normal food vials to agar-only vials. Affixed to the vials was a rubber stopper with one glass pipet (World Precision Instruments, #1B150F-4), initially empty. Animals were starved in this chamber overnight, also allowing for habituation to the glass pipet. At ZT 1, the empty glass pipet was replaced with a pipet filled with 2% sucrose solution, dyed blue (FD&C Blue #1, Fisher #S06652) and topped with a drop of mineral oil to prevent evaporation. Meniscus movement was measured over 6 hours. Both the habituation and experimental periods were conducted in a 25C humidity-controlled incubator.

### Starvation resistance assay

Starvation resistance assay was conducted in using DAM system as previously described. Flies were loaded into the single beam DAM system as previously described for sleep assays, except DAM tubes were filled with 2% agar-only food. Data was collected until all animals were deceased, roughly 4 days.

### qPCR

20 female adult brains per sample were dissected on 1X phosphate-buffered saline, with 3 biological replicates per genotype. RNA was then extracted using the RNeasy Plus Mini Kit (Qiagen, #74136). Primers were designed against *beat-Ia A* and *B* isoforms, and neuronal tubulin was used as a housekeeping gene. RNA was amplified using SsoAdvanced SYBR Green Supermix and amplification was measured using a CFX Opus 96 Real-Time PCR System. Relative changes in beat mRNA expression were quantified using the 2^−ΔΔCt^ method.

Primers against *beat-Ia AB*
Forward: GTCCTCGAACGAGAGCCAAG
Reverse: GCATTGTGGGGCGTTTCAATAA

### Immunohistochemistry

Fly brains were dissected in 1X phosphate-buffered saline (PBS) and fixed in 4% paraformaldehyde (PFA) for 15 min at room temperature. Samples were then washed three times for 15 min each in PBS with 0.1% Triton X-100 (PBST) and then blocked in PBST with 5% goat serum for 1 hr at room temperature. Samples were then incubated with primary antibody in PBST with 5% goat serum at 4C overnight. Samples were then washed three times for 15 min each in PBST and incubated with secondary antibody in PBST with 5% goat serum at 4C overnight. Samples were then washed three times for 15 min each in PBST, cleared in 50% glycerol, and mounted in VECTASHIELD. The following primary antibodies were used: mouse anti-side (1:100, DSHB), rabbit anti-GFP (1:500, Invitrogen), mouse anti-GFP (1:500, Fisher), rabbit anti-NPF (1:250, RayBioTech), mouse anti-nc82 (1:100, DSHB). The following secondary antibodies were used (all at 1:250, Invitrogen): goat anti-mouse 488, donkey anti-mouse 594, goat anti-rabbit 488, and goat anti-rabbit 594.

### Imaging

Microscopy images were taken using a Leica TCS SP8 confocal microscope. Unless otherwise specified, maximum projection images were generated of all Z-slices, with images taken in 0.5 µM steps. If necessary for whole-brain visualization, images were stitched using GIMP.

### Syt-EGFP puncta analysis

Maximum projection images of Syt-EGFP samples were generated and analysis was conducted in ImageJ. Cell bodies were selected and deleted to avoid quantification of extraneous signal. An ROI of the entire sample was defined, using nc82 as an anatomical stain. The EGFP channel was isolated from the merged image and converted to a binary image using ImageJ’s Intermodes dark no-reset threshold. The Analyze Particles function was then used to count all objects larger than 1.50 pixels^2^.

### Statistical analysis

All statistical analyses were performed using GraphPad Prism. Details on sample sizes and statistical tests are denoted in figure legends.

## Acknowledgments

We thank David Raizen, Amita Sehgal, members of the Kayser Lab, members of the Raizen Lab, and members of the Penn Chronobiology and Sleep Institute for helpful discussions and input.

## Funding

NIH T32GM008076 (KM)

NIH T32HL007953 (KM)

NIH T32HL007713 (AZ)

NIH R35HG011959 (AC)

Fulbright Visiting Scholar Program – Postdoctoral Grant (FY-2017-TR-PD-07) (FDB)

NIH P01HL160471 (AIP)

NIH R01HL143790 (SFG)

Daniel B. Burke Endowed Chair for Diabetes Research (SFAG)

NIH DP2NS111996 (MSK)

NIH R01NS120979 (MSK)

Linda Pechenik Montague Award (MSK)

Burroughs Wellcome Career Award for Medical Scientists (MSK)

## Author contributions

Conceptualization: KM, AZ, AC, AI, SFAG MSK

Investigation: KM, AZ, AC, FDB, HK, EADV

Writing – Original Draft: KM, MSK

Writing – Review and Editing: All authors

Project Supervision and Funding: MSK

## Competing interests

Authors declare that they have no competing interests.

## Data and materials availability

All data needed to evaluate the conclusions in the paper are present in the paper and/or the Supplementary Materials.

## Supplemental Figures and Tables

**Supplemental Fig. 1.**
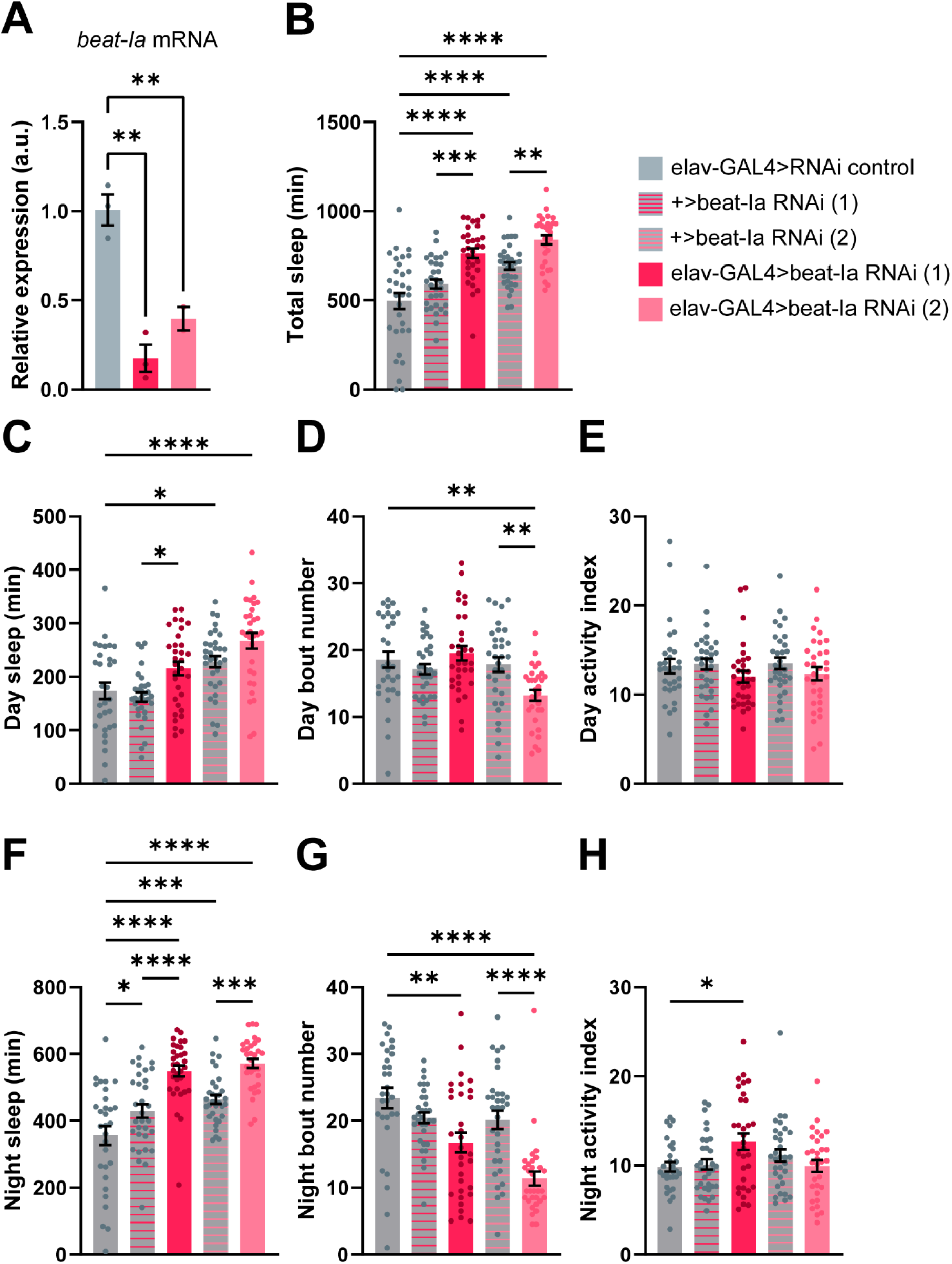
Confirmation of *beat-Ia* knockdown phenotype with qPCR, high resolution sleep measurement. (A) qPCR confirms effective *beat-Ia* knockdown from two independent RNAi. Relative expression was calculated using the 2-ΔΔCT method from 2-3 biological replicates per genotype, each with 3 technical replicates. One-way ANOVA with Dunnett’s multiple comparisons tests comparing RNAi genotypes to control. n, from left to right = 3, 3, 2. (B-H) Sleep measures from multibeam replicates of pan-neuronal beat-Ia knockdown animals. (B) Total sleep. (C-E) Day sleep measures. (F-H) Night sleep measures. One-way ANOVA with Šídák’s multiple comparisons tests comparing experimental genotypes to their respective controls. n, from left to right = 30, 32, 32, 32, 31.

**Supplemental Fig. 2.**
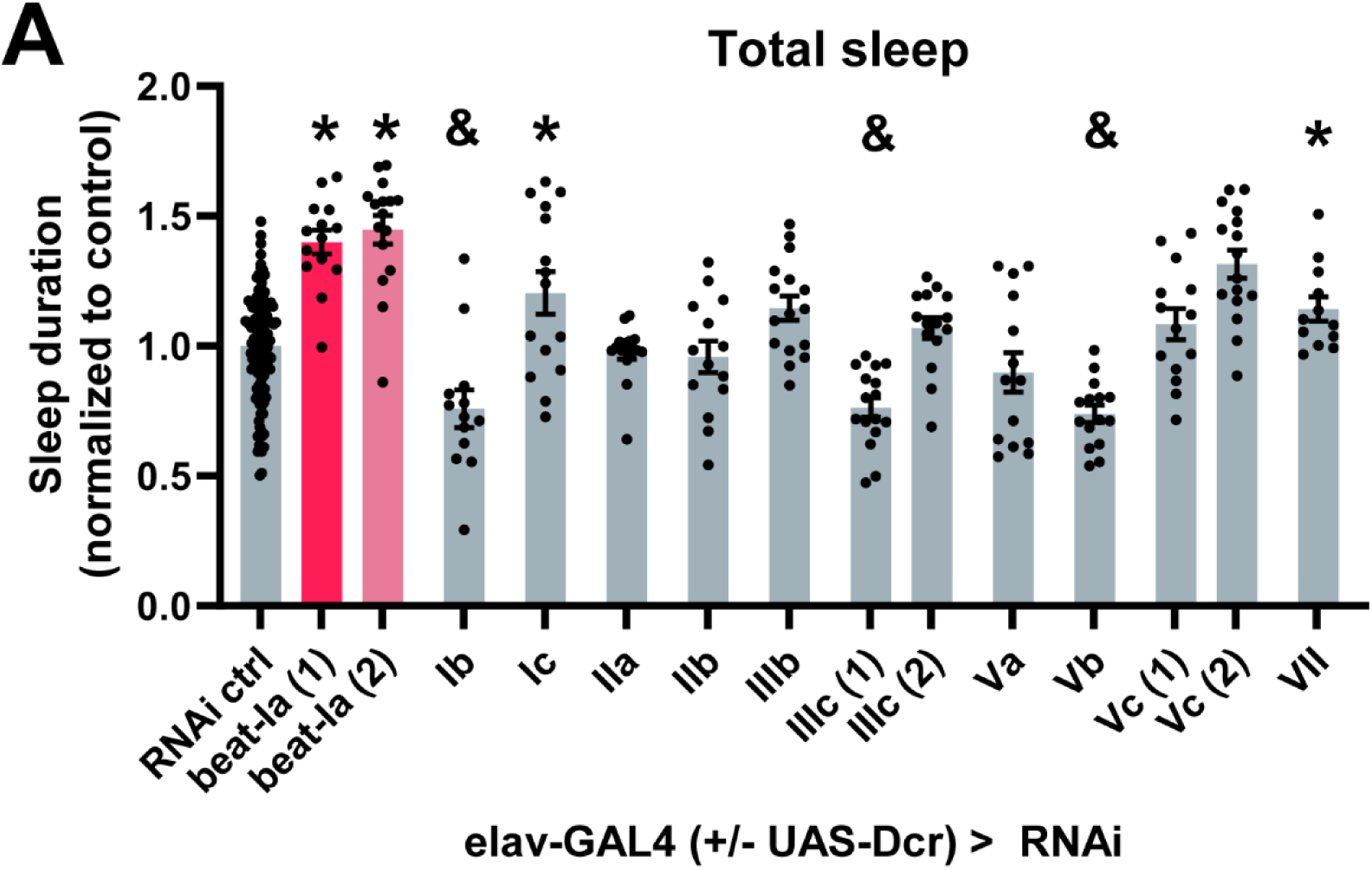
Knockdown of other *beat* isoforms results in sleep phenotypes. (A) Normalized total sleep for animals expressing RNAi against members of the beaten path family of adhesion molecules. * indicates significant increase in sleep, & indicates significant decrease in sleep compared to RNAi control. One-way ANOVA with Dunnett’s multiple comparisons tests comparing experimental genotypes to RNAi control. n, from left to right = 124 (pooled from all experiments), 14, 16, 13, 15, 16, 14, 16, 16, 15, 14, 15, 14, 16, 12.

**Supplemental Fig. 3.**
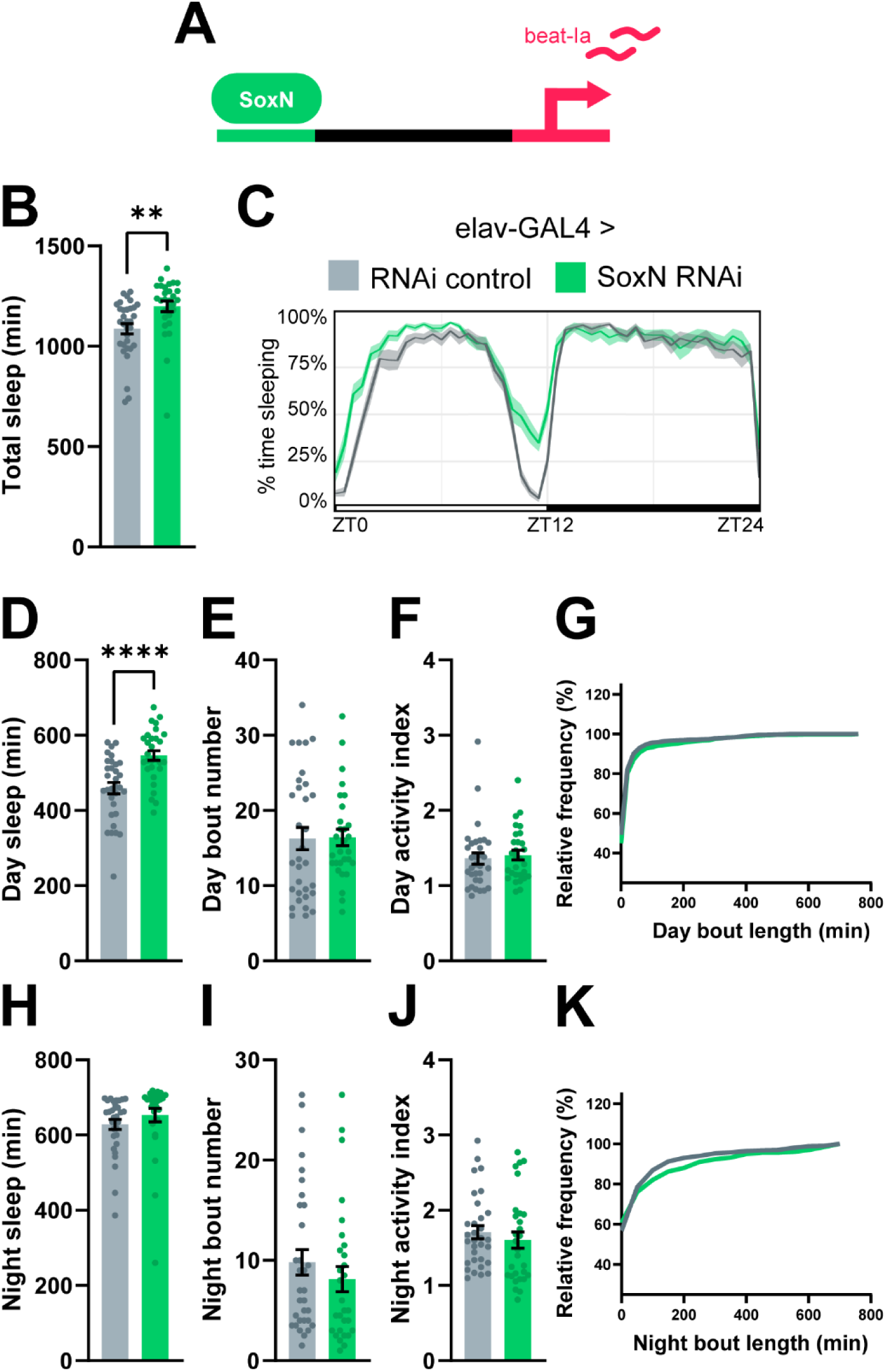
Knockdown of *SoxN*, a transcription factor upstream of beat-Ia, also increases sleep. (A) Schematic of *SoxN*>*beat-Ia* signaling pathway. (B) Total sleep. (C) Sleep trace. (D-G) Day sleep measures. (H-K) Night sleep measures. Unpaired t test. n, left to right = 32, 29.

**Supplemental Fig. 4.**
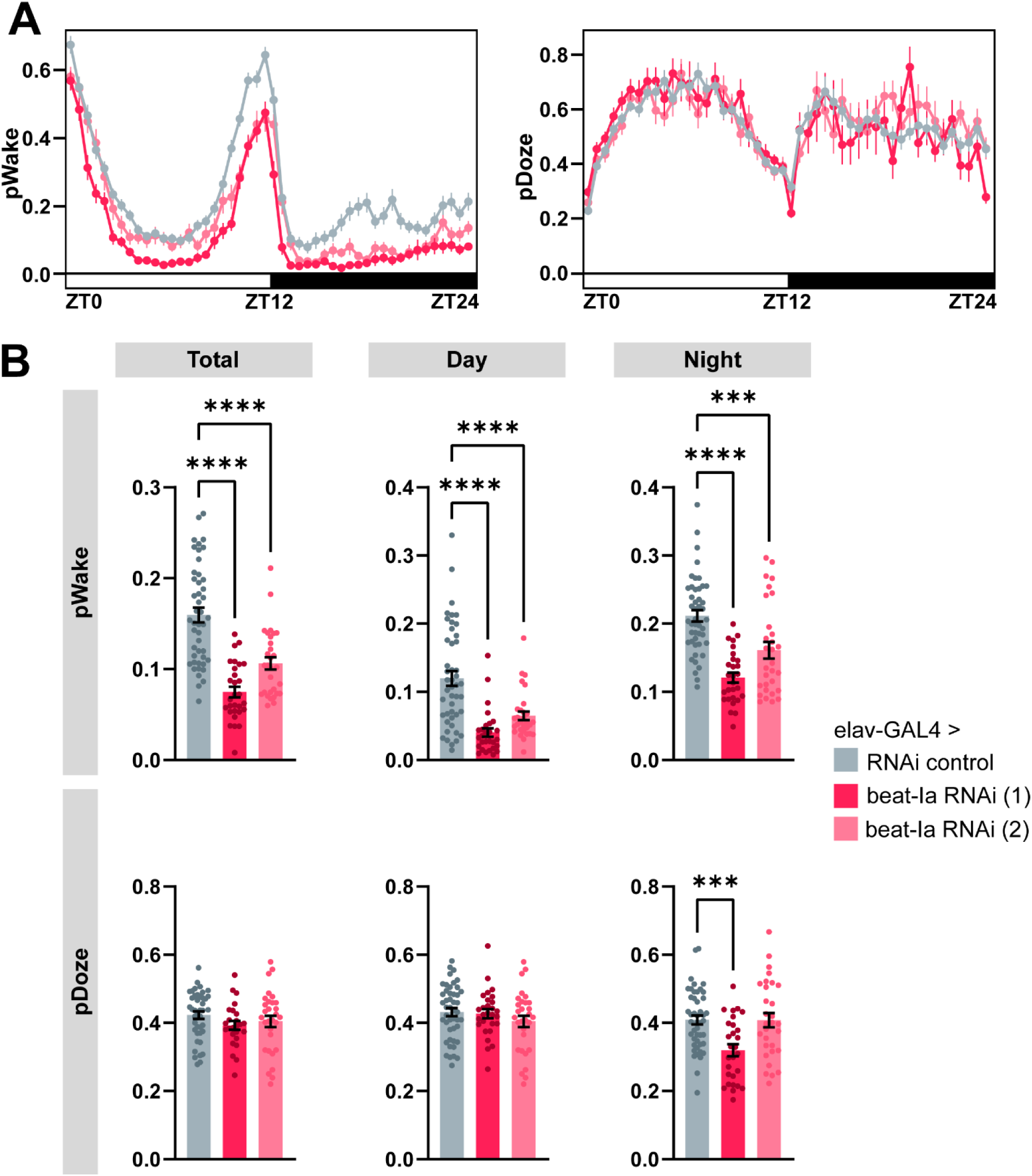
*beat-Ia* knockdown animals are less likely to wake from a sleep state. (A) Probability of transitioning from a sleep state to a wake state (pWake) or from a wake state to a sleep state (pDoze) in rolling 30min bins across 24hrs. (B) Average pWake/pDoze across the day, night, and total 24hrs. One-way ANOVA with Dunnett’s multiple comparisons tests comparing experimental genotypes to RNAi control. n, from left to right = 46, 29, 30.

**Supplemental Fig. 5.**
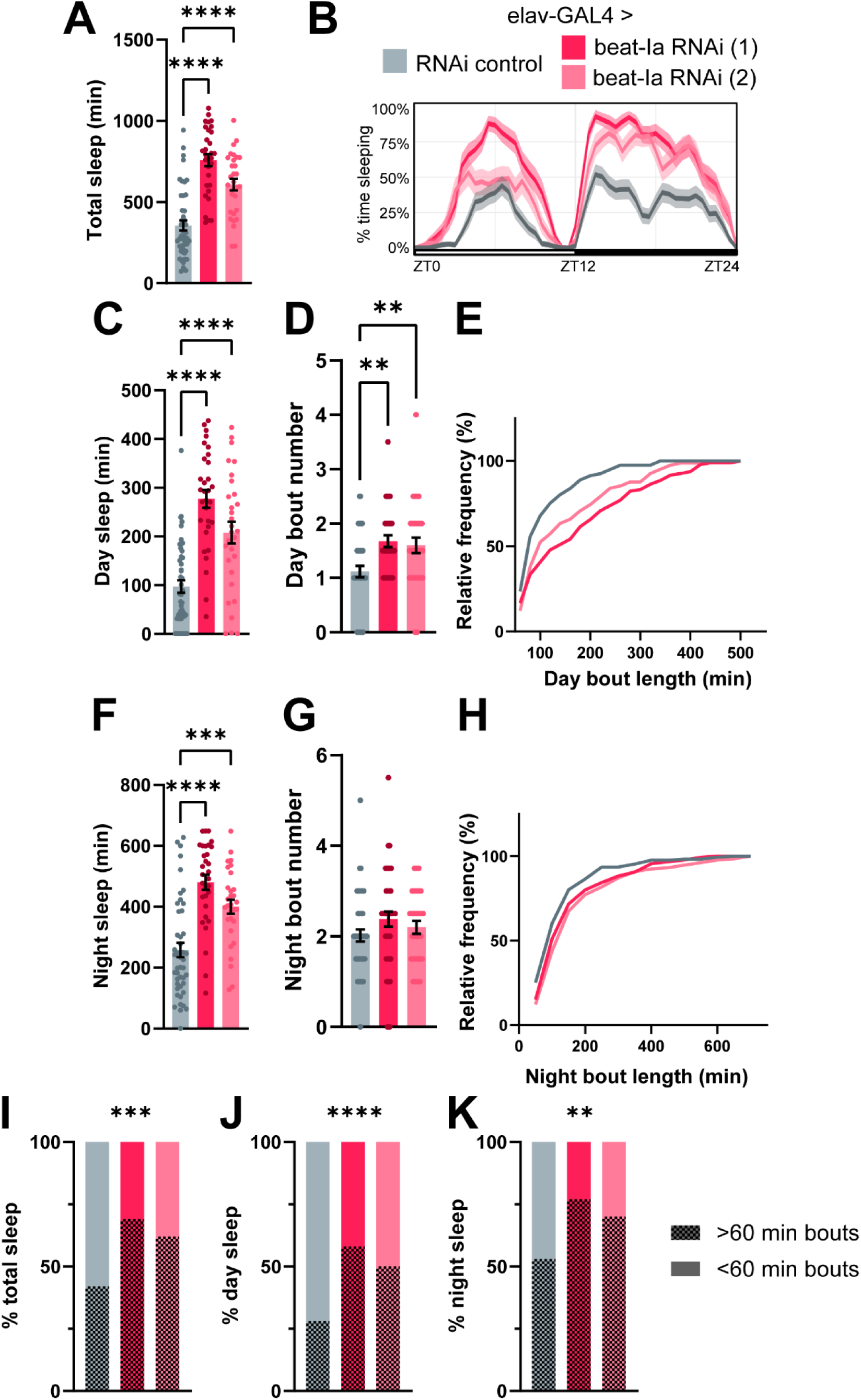
*beat-Ia* knockdown animals spend more time in long sleep bouts than controls. Sleep measures from only sleep bouts ≥ 60 min. (A) Total sleep. (B) Sleep trace. (D-E) Day sleep measures. (F-H) Night sleep measures. One-way ANOVA with Dunnett’s multiple comparisons tests comparing experimental genotypes to RNAi control. n, from left to right = 46, 31, 30. (I-K) Average percent time spent in long sleep bouts vs. bouts shorter than 60 min. Fisher’s exact test.

**Supplemental Fig. 6.**
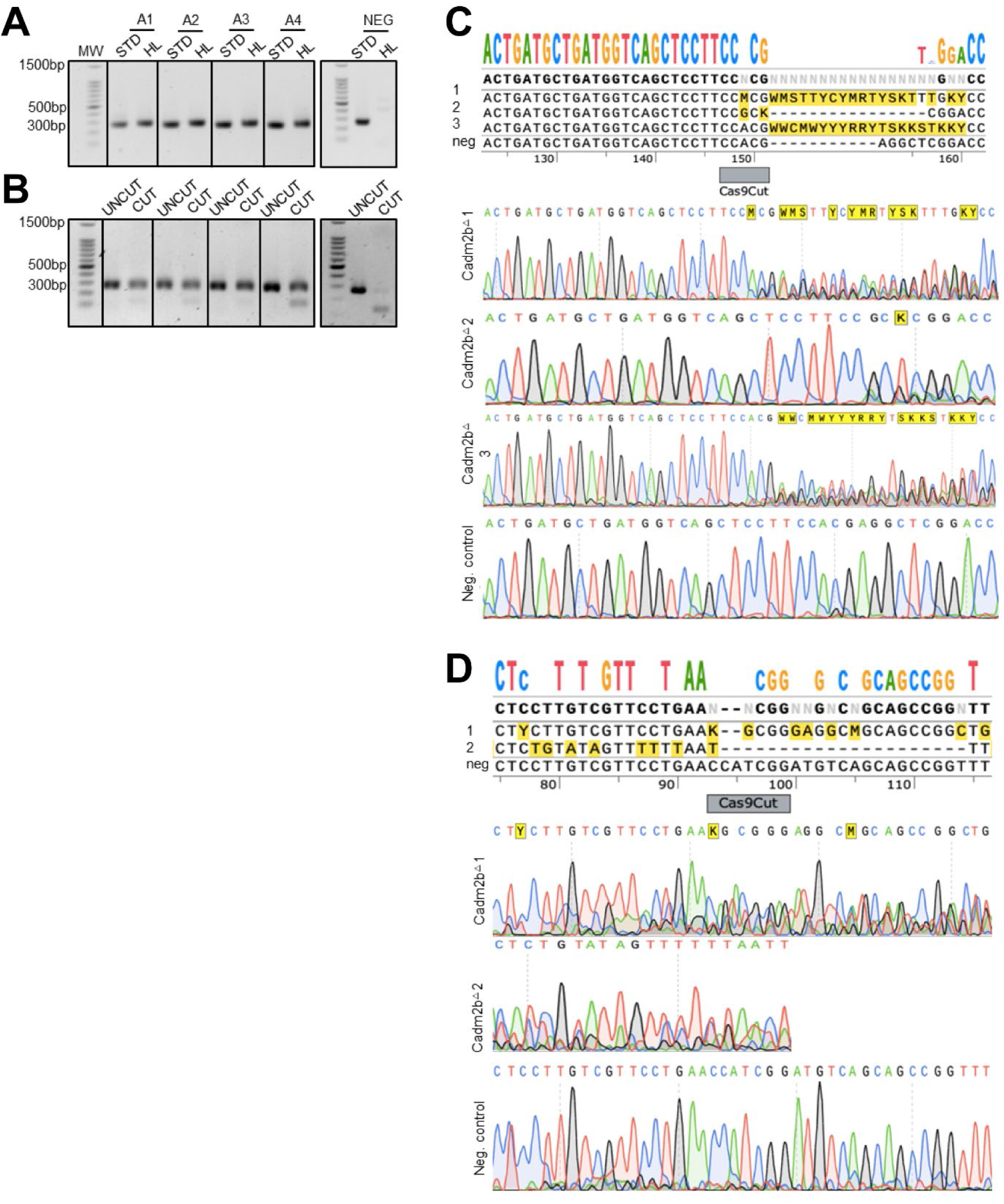
On-target mutation confirmation of guide RNAs for zebrafish F0 CRISPR lines. Following the sleep assay, fish are genotyped using headloop PCR (A) for guideRNA 1 and restriction digest PCR (B) for guideRNA 2. (A-B) PCR results showing 4 crispant fish (A1-4) and one negative control (NEG). (A) Headloop PCR for guideRNA 1 confirms mutants with band present (mutation presence inhibits suppression of amplification) using both standard (STD) and headloop (HL) primers. In the wild-type sequence, the headloop primer forms a small hairpin that leads to suppression of amplification. Negative control shows suppression of the wild-type DNA, so no band is present in the HL column. (B) Restriction digest PCR for guideRNA 2 using the same fish from (A). The guideRNA is designed so the cut site overlaps a restriction enzyme site. If a mutation is present, the sequence will not be cut by the enzyme and will show a band of the same size as the uncut sequence. Negative control shows restriction enzyme is able to cut the sequence indicating a wild-type sequence. (C) Representative electropherograms and sequence alignment of the on-target mutation for 3 crispants and 1 negative control. Reverse sequence is shown. Guide RNA with respect to the reverse strand is CCACGAGGCTCGGACCAGTTCAA; therefore, the Cas9 predicted cut site would be 3bp upstream of the PAM (CCA). (D) Representative electropherograms and sequence alignment of the on-target mutation for 2 crispants and 1 negative control. Reverse sequence is shown. Guide RNA with respect to the reverse strand is CCTTGTCGTTCCTGAACCATCGG ; therefore, the Cas9 predicted cut site would be 3bp upstream of the PAM (CGG). Note, because this guide RNA is downstream (exon 4) of guide RNA1 (exon 3), different frame shifts have occurred due to in/dels leaving the sequence highly mixed.

**Supplemental Fig. 7.**
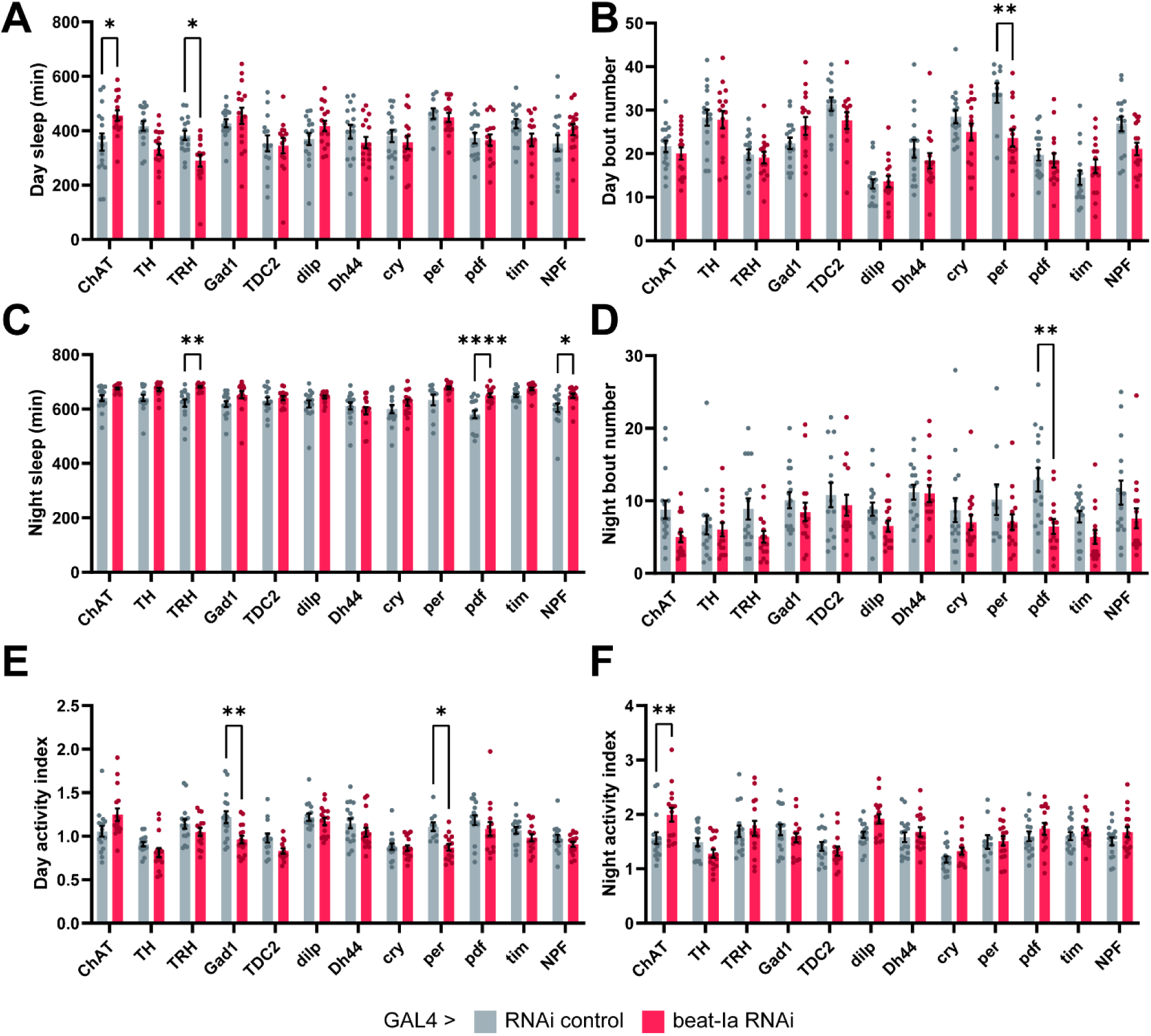
Targeted spatial *beat-Ia* knockdown screen. Sleep measures from animals expressing *beat-Ia* RNAi in populations of sleep- and/or circadian-regulatory cells. (A-B) Day sleep measures. (C-D) Night sleep measures. (E-F) Activity indices. Two-way ANOVA with Šídák’s multiple comparisons tests comparing each experimental genotype to its control. n = 16 except for TRH-GAL4>*beat-Ia* RNAi (15), TDC2-GAL4>RNAi control (14), TDC2-GAL4>*beat-Ia* RNAi (14), per-GAL4>RNAi control (10), per-GAL4>*beat-Ia* RNAi (15), and pdf-GAL4>*beat-Ia* RNAi (15).

**Supplemental Fig. 8.**
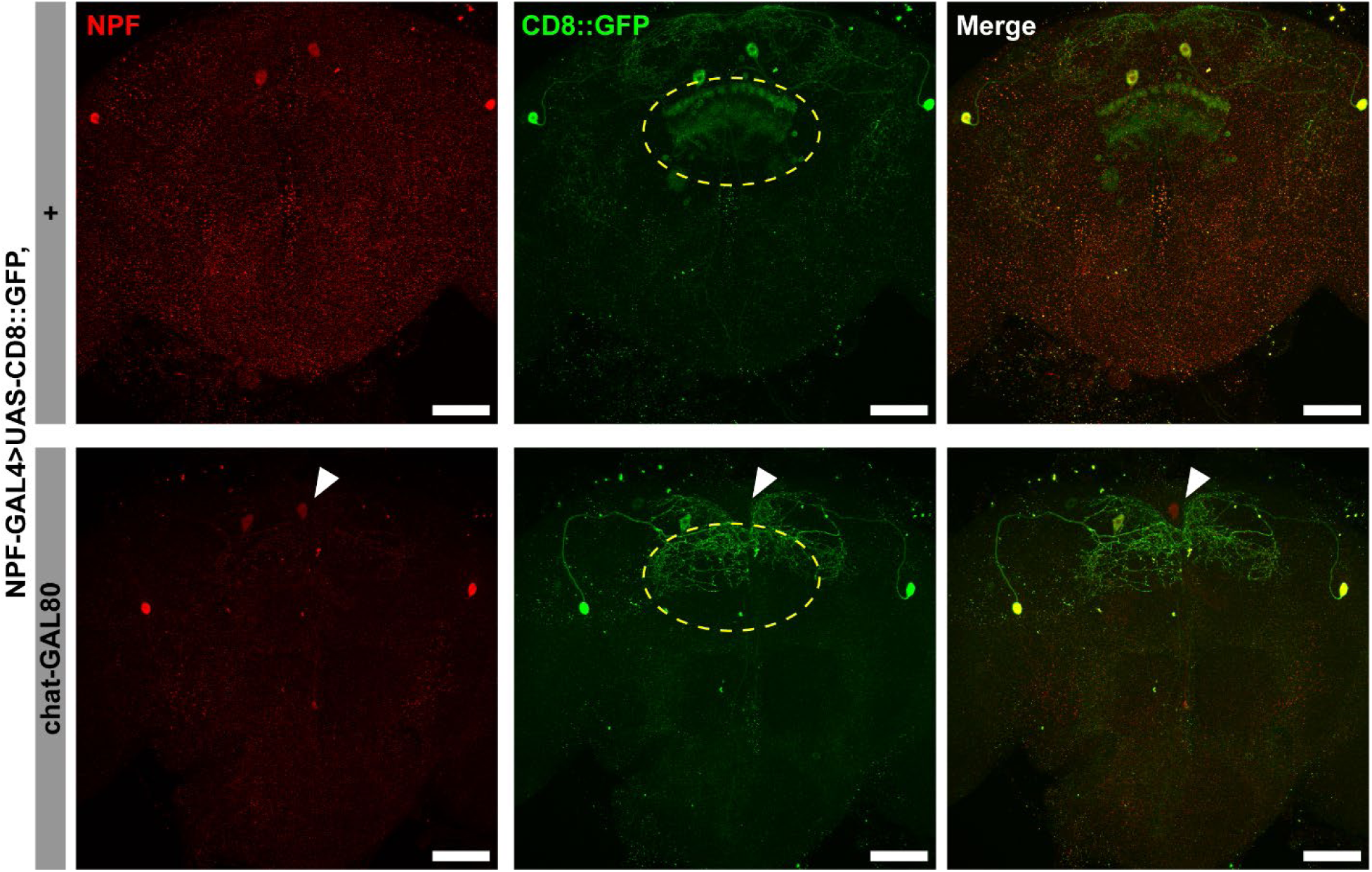
Some NPF cells co-express choline acetyltransferase (ChAT). Maximum projection images of brains from flies expressing UAS-CD8::GFP in NPF cells, with or without chat-GAL80. chat-GAL80 appears to inhibit UAS-CD8::GFP expression in some large descending NPF cells (white arrow) and a cluster of NPF cells in the fan-body (yellow dotted oval). Scale bar = 50µM.

**Supplemental Fig. 9.**
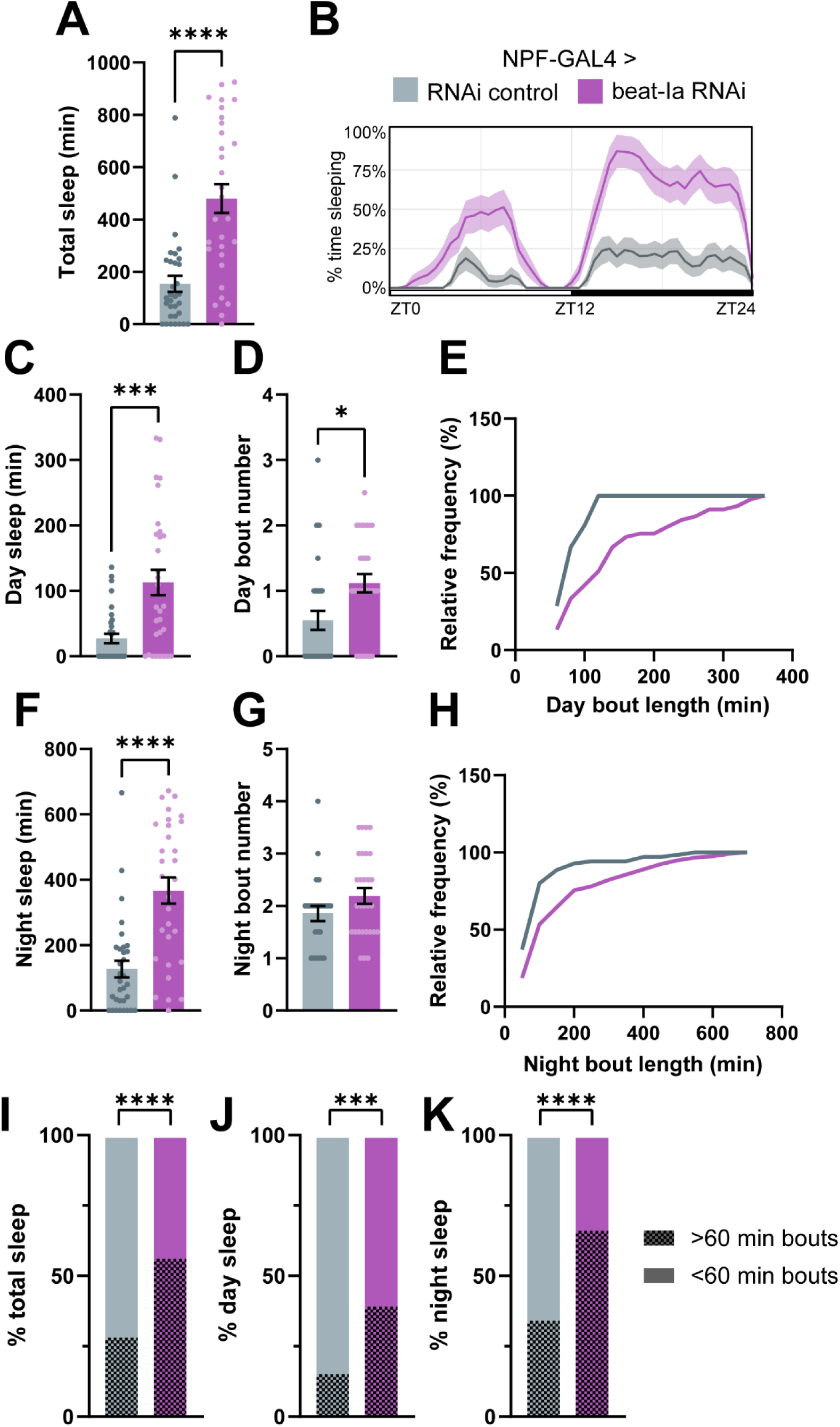
NPF *beat-Ia* knockdown animals spend more time in long sleep bouts. Sleep measures from only sleep bouts ≥ 60 min. (A) Total sleep. (B) Sleep trace. (C-E) Day sleep measures. (F-H) Night sleep measures. Welch’s t test. n, left to right = 32, 30. (I-K) Average percent time spent in long sleep bouts vs. bouts shorter than 60 min. Fisher’s exact test.

**Supplemental Fig. 10.**
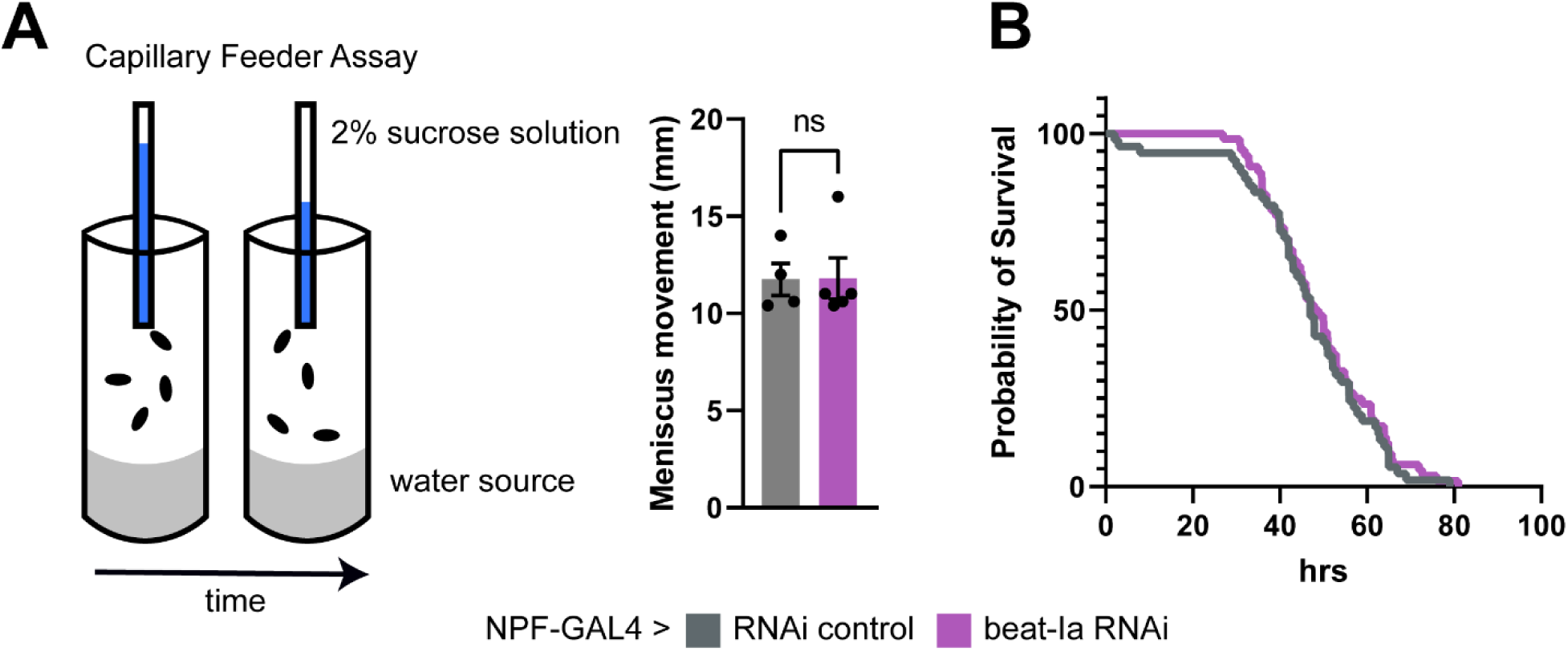
*beat-Ia* knockdown in NPF cells does not affect feeding or resistance to starvation. (A) Schematic of Capillary Feeder Assay (CAFÉ). Feeding is measured by movement of the meniscus of a dyed sucrose solution in a glass capillary. (B) *beat-Ia* knockdown and control animals feed similarly in the CAFÉ assay. Each data point represents one vial containing 10 flies. Unpaired t test. n, left to right = 4, 5. (B) No difference in resistance to starvation between *beat-Ia* knockdown and control animals. Log-rank (Mantel-Cox) test. n = 54 (NPF-GAL4>RNAi control), 64 (NPF-GAL4>*beat-Ia* RNAi).

**Supplemental Fig. 11.**
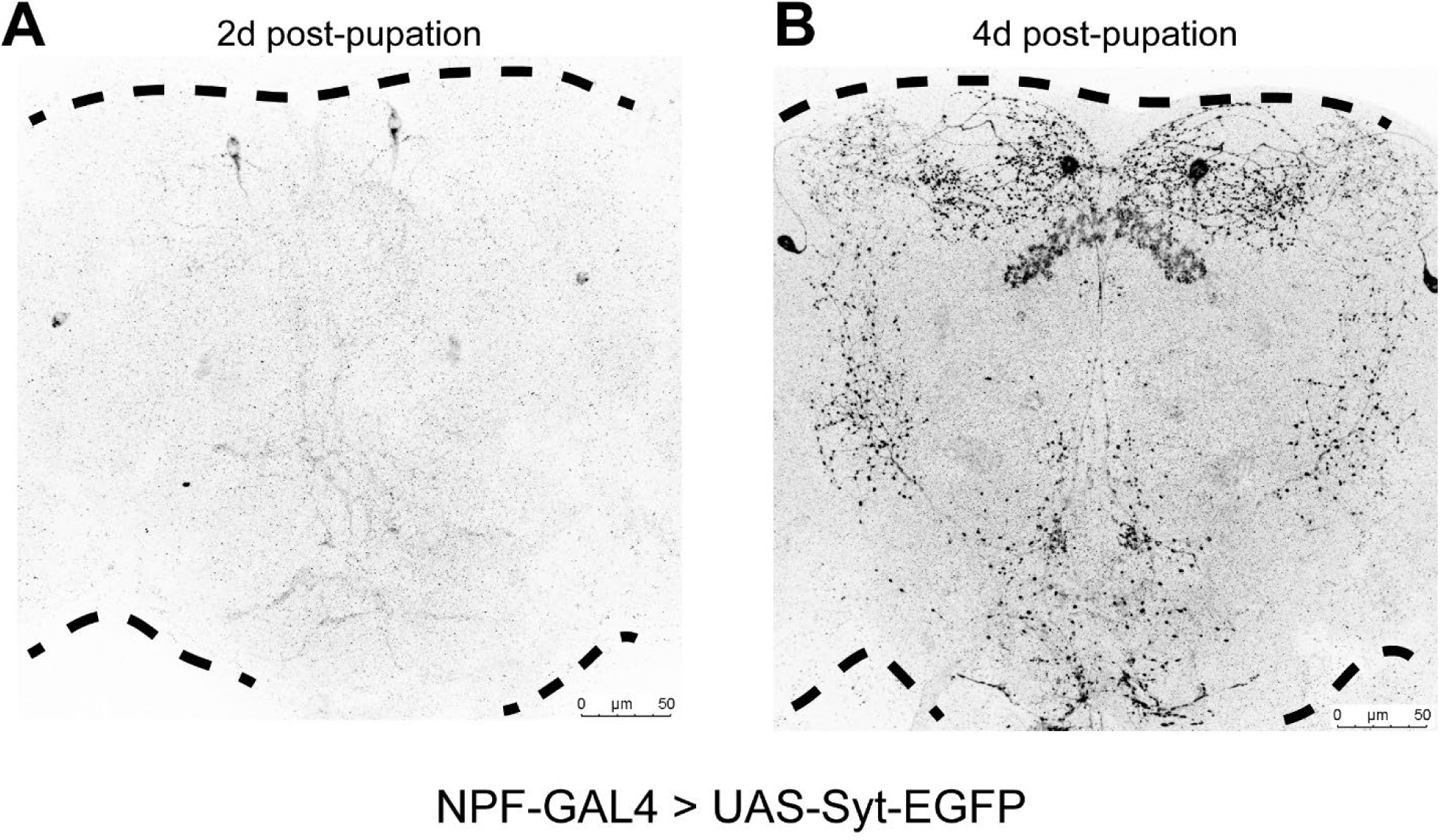
NPF innervates the SEZ in the mid-pupal stages. Maximum projection images of brains from NPF-GAL4>UAS-Syt-EGFP animals at 2d (A) and 4d (B) post-pupation. Scale bar = 50µM.

**Supplemental Fig. 12.**
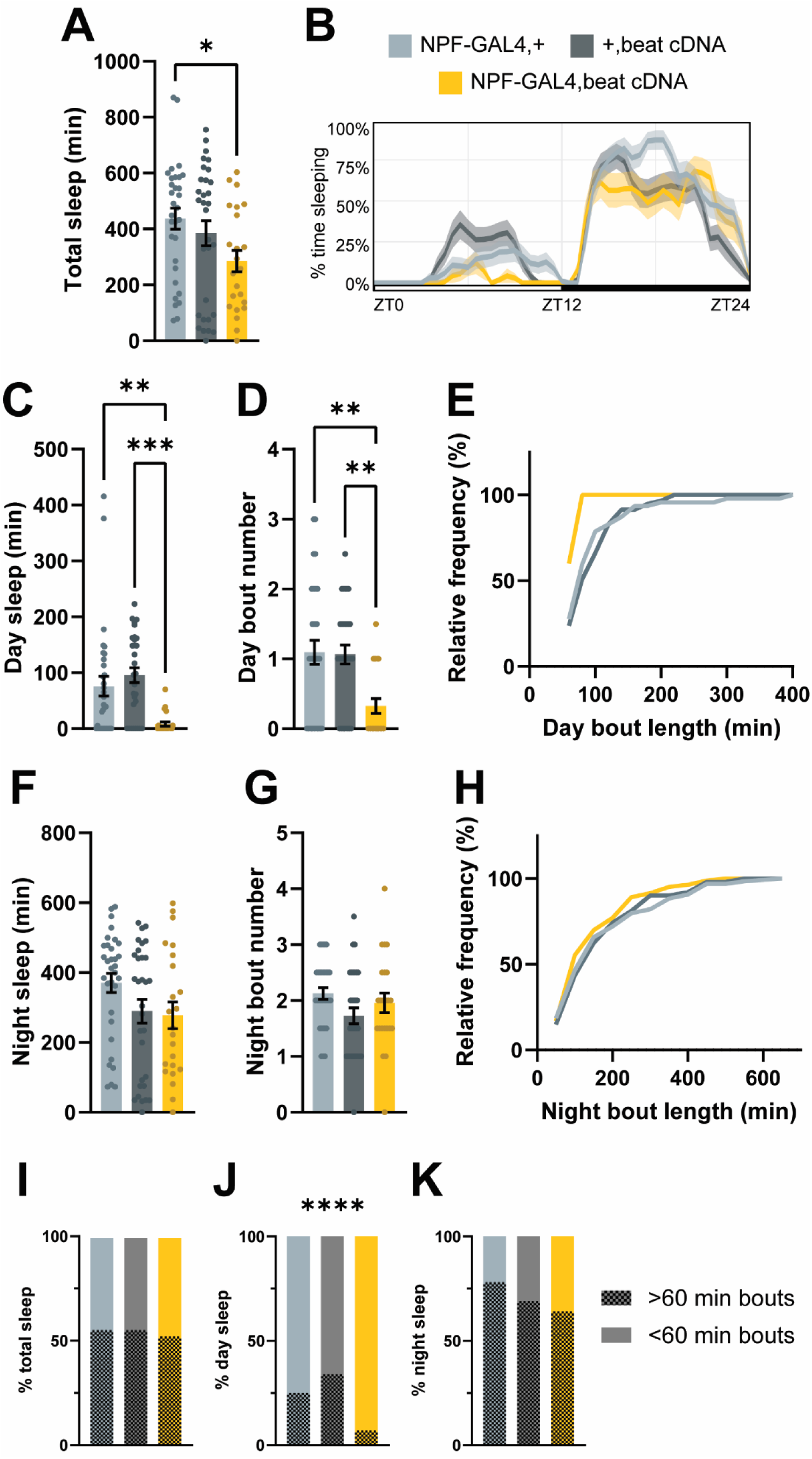
*beat-Ia* overexpression in NPF cells reduces the time spent in long sleep bouts. Sleep measures from only sleep bouts ≥ 60 min. (A) Total sleep. (B) Sleep trace. (C-E) Day sleep measures. (F-H) Night sleep measures. One-way ANOVA with Tukey’s multiple comparisons tests comparing experimental genotype to controls. n, from left to right = 32, 31, 23. (I-K) Average percent time spent in long sleep bouts vs. bouts shorter than 60 min. Fisher’s exact test.

**Supplemental Fig. 13.**
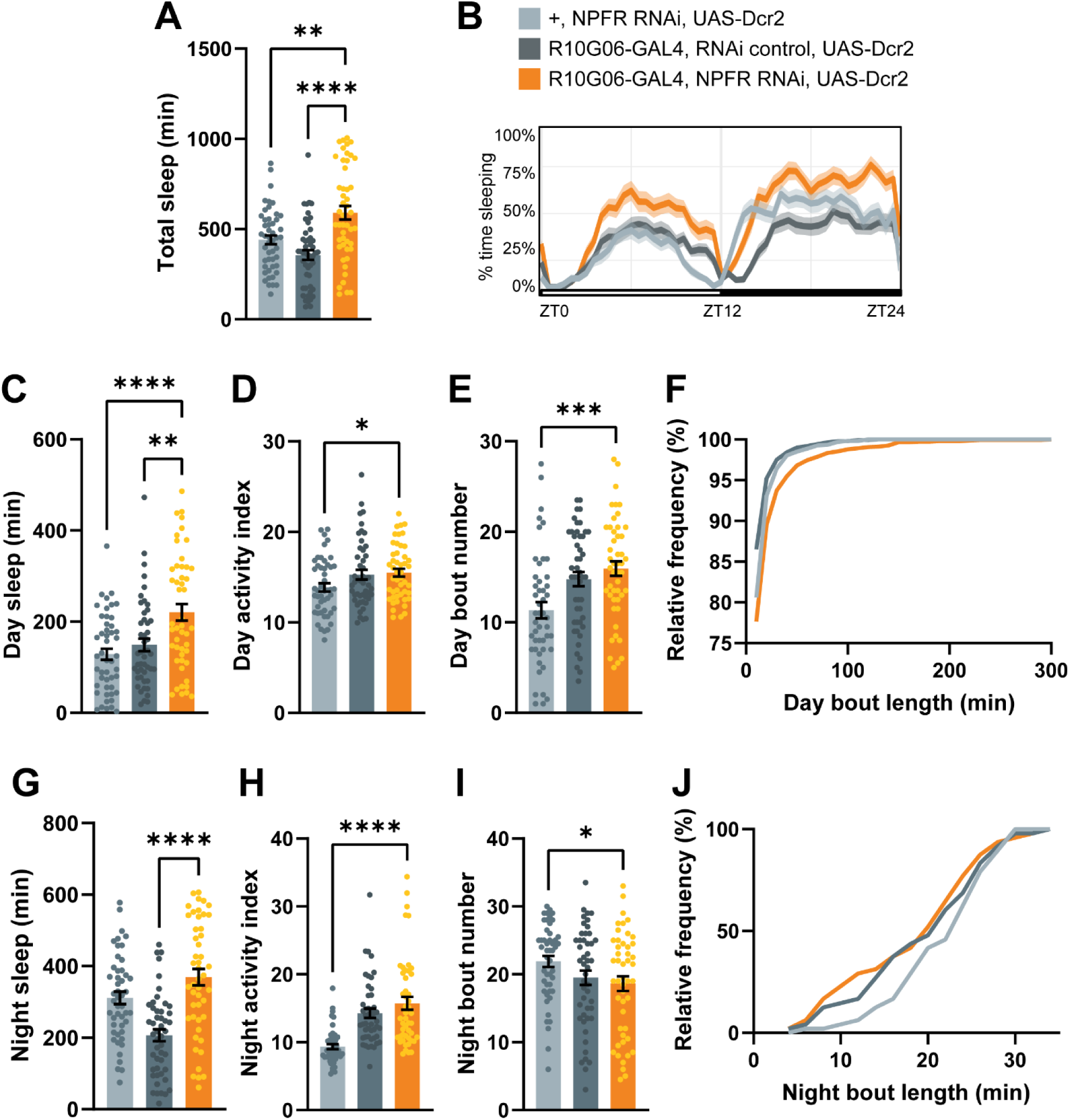
Knockdown of *NPFR* in 10G06-GAL4-labeled cells increases sleep. (A) Total sleep. (B) Sleep trace. (C-F) Day sleep measures. (G-J) Night sleep measures. One-way ANOVA with Dunnett’s multiple comparisons tests comparing experimental genotype to controls. n = 48 for all genotypes.

**Supplemental Fig. 14.**
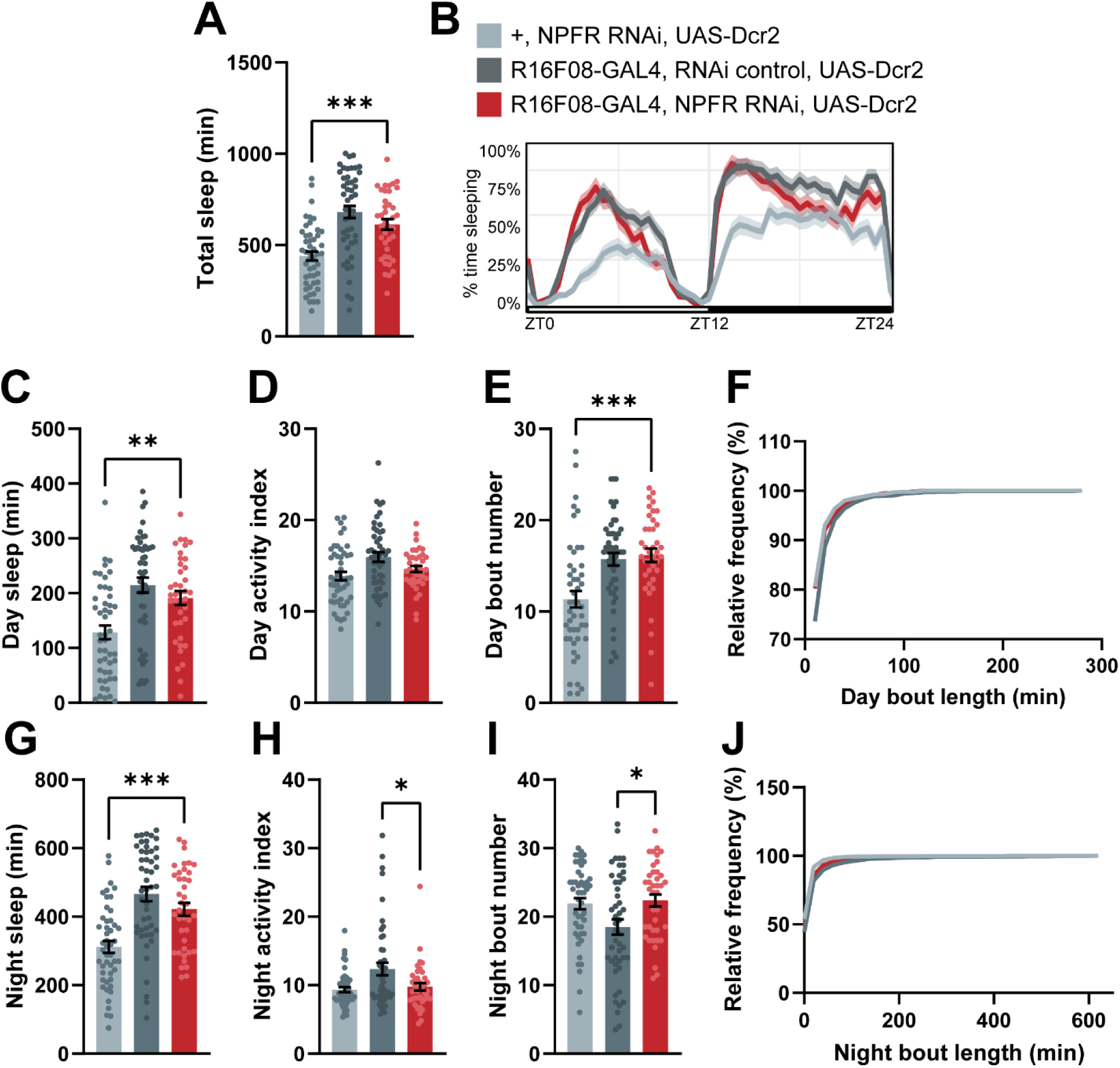
Knockdown of *NPFR* in 16F08-GAL4-labeled cells does not increase sleep upon replication. (A) Total sleep. (B) Sleep trace. (C-F) Day sleep measures. (G-J) Night sleep measures. One-way ANOVA with Dunnett’s multiple comparisons tests comparing experimental genotype to controls. n, from left to right = 39, 48, 48.

**Supplemental Fig. 15.**
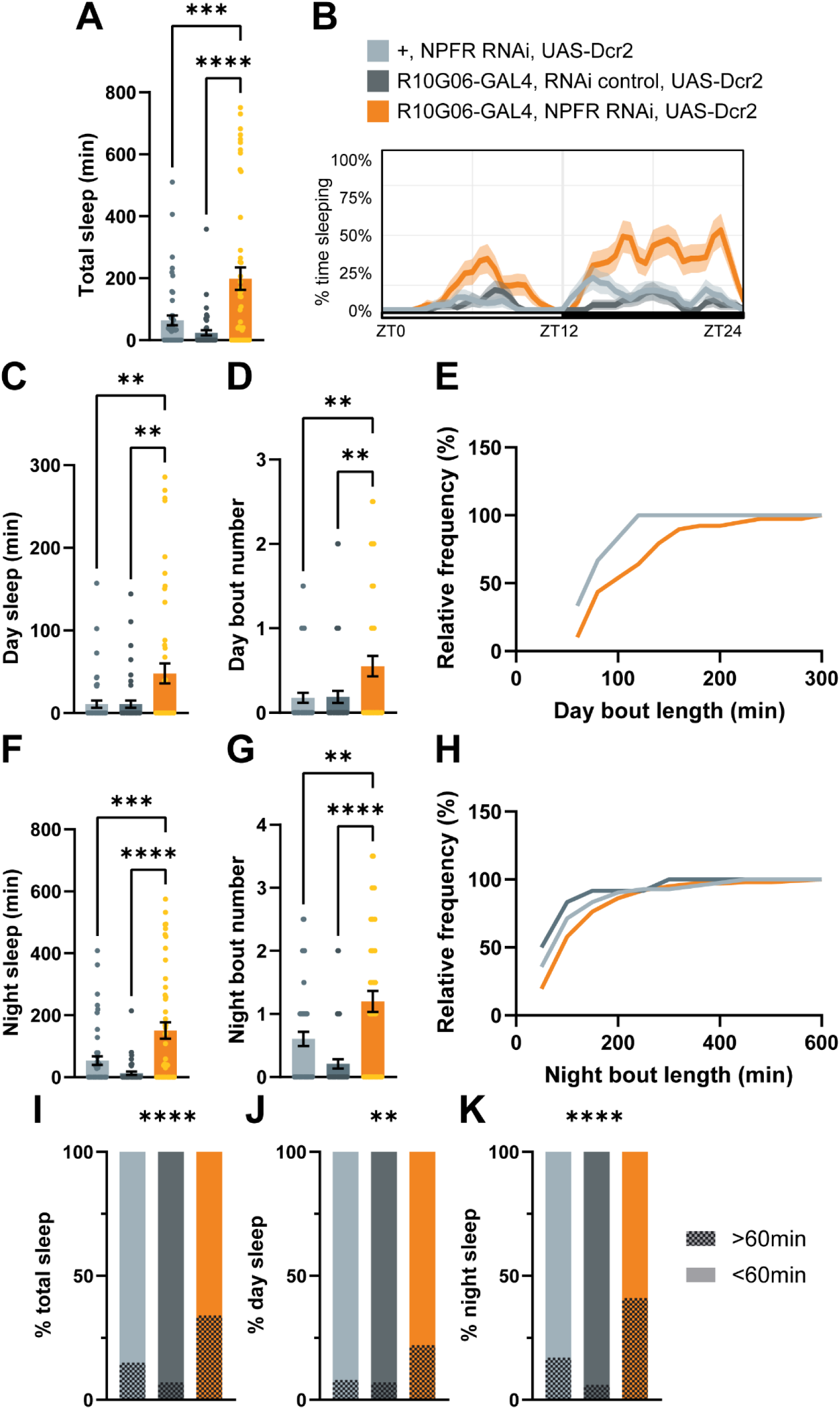
10G06-GAL4>*NPFR* knockdown animals spend more time in long sleep bouts. Sleep measures from only sleep bouts ≥ 60 min. (A) Total sleep. (B) Sleep trace. (C-E) Day sleep measures. (F-H) Night sleep measures. (I-K) Average percent time spent in long sleep bouts vs. bouts shorter than 60 min. For (E), +,*NPFR* RNAi,UAS-Dcr2 and R10G06-GAL4,RNAi control,UAS-Dcr2 lines are identical so only one control is visible. One-way ANOVA with Dunnett’s multiple comparisons tests comparing experimental genotype to controls. n = 48 for all genotypes.

**Supplemental Fig. 16.**
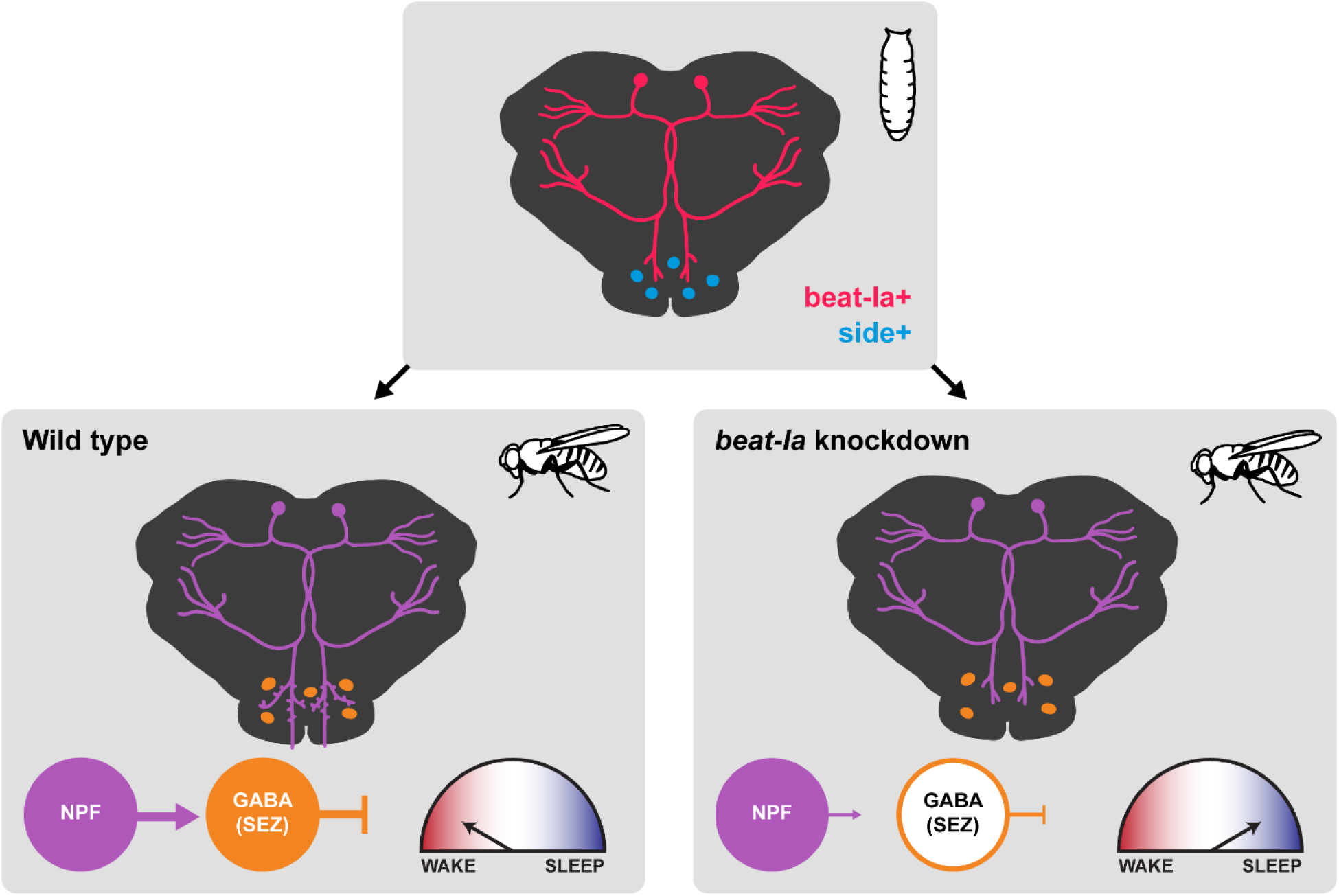
*beat-Ia* acts in development to pattern an NPF>SEZ sleep-regulatory circuit. Schematic illustrating a working model of *beat-Ia*’s role in patterning adult sleep circuits. Top panel: During mid- to late-pupal development, *beat-Ia*-expressing NPF cells are properly connected to the SEZ in a manner coordinated by *side*-expressing cells. Bottom panel, left: In wild-type animals, NPF cells synapse onto GABAergic SEZ cells, which inhibit downstream sleep-regulatory outputs to maintain wakefulness. Bottom panel, right: *beat-Ia* knockdown in NPF cells results in a loss of synaptic density in the SEZ, reducing excitatory input onto GABAergic cells, and impaired ability to maintain wakefulness due to disinhibition of downstream sleep-regulatory outputs.

**Supplemental Fig. 17.**
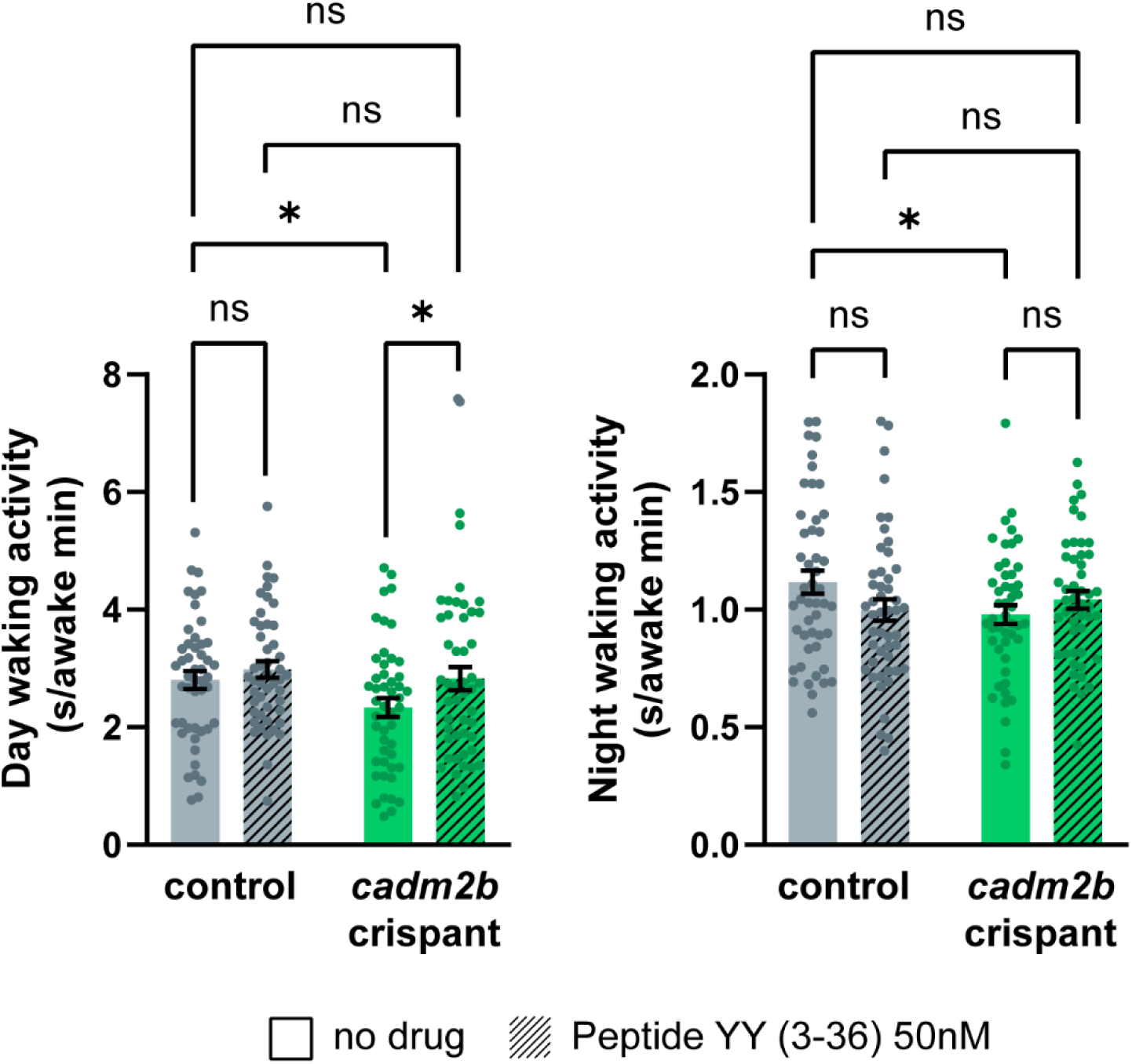
Waking activity in control and *cadm2b* crispant zebrafish with and without NPY receptor agonism. Activity during day wake bouts (left) and night wake bouts (right). Two-way ANOVA with Uncorrected Fisher’s LSD post-hoc tests comparing within genotypes no drug to Peptide YY, within drug conditions negative control siblings to *cadm2b* crispants, and negative control siblings with no drug to *cadm2b* crispant on Peptide YY. n = 48 for all genotypes.

**Supplemental Data 1. GWAS loci and RNAi screen data.**

(A) All sleep propensity SNPs identified from sleep and hypersomnia GWAS and the genes in their respective TADs. (B) All unique coding genes identified in (A), excluding non-coding genes. (C) Sleep measures, normalized to an RNAi control, for each RNAi driven by a pan-neuronal driver.

**Supplemental Data 2. GAL4s matched to putative NPF>SEZ cells.**

(A) Full list of GAL4s identified as potential matches to FlyWire-identified NPF>SEZ cells. (B) List of GAL4s screened for sleep phenotypes and the cells to which they were matched.

## Notes

### Competing Interest Statement

The authors have declared no competing interest.

